# Effect of insulin insufficiency on ultrastructure and function in skeletal muscle

**DOI:** 10.1101/2022.12.18.520900

**Authors:** Chantal Kopecky, Michael Haug, Barbara Reischl, Nandan Deshpande, Bikash Manandhar, Thomas W. King, Victoria Lee, Marc R Wilkins, Margaret Morris, Patsie Polly, Oliver Friedrich, Kerry-Anne Rye, Blake J Cochran

**Affiliations:** School of Biomedical Sciences, Faculty of Medicine & Health, UNSW Sydney, Sydney, Australia; Institute of Medical Biotechnology, Department of Chemical and Biological Engineering, Friedrich-Alexander University Erlangen-Nürnberg, Erlangen, Germany; Sydney Informatics Hub, University of Sydney, Sydney, Australia; Systems Biology Initiative, Faculty of Science, UNSW Sydney, Sydney, Australia

**Keywords:** Insulin, skeletal muscle, muscle structure, muscle function

## Abstract

**Background:** Decreased insulin availability and high blood glucose levels, the hallmark features of poorly controlled diabetes, drive disease progression and are associated with decreased skeletal muscle mass. We have shown that mice with β-cell dysfunction and normal insulin sensitivity have decreased skeletal muscle mass. This project asks how insulin deficiency impacts on the structure and function of the remaining skeletal muscle in these animals.

**Methods:** Skeletal muscle function was determined by measuring exercise capacity and specific muscle strength prior to and after insulin supplementation for 28 days in 12-week-old mice with conditional β-cell deletion of the ATP binding cassette transporters ABCA1 and ABCG1 (β-DKO mice). *Abca1* and *Abcg1* floxed (fl/fl) mice were used as controls. RNAseq was used to quantify changes in transcripts in soleus and extensor digitorum longus muscles. Skeletal muscle and mitochondrial morphology were assessed by transmission electron microscopy. Myofibrillar Ca^2+^ sensitivity and maximum isometric single muscle fibre force were assessed using *MyoRobot* biomechatronics technology.

**Results:** RNA transcripts were significantly altered in β-DKO mice compared to fl/fl controls (32 in extensor digitorum longus and 412 in soleus). Exercise capacity and muscle strength were significantly decreased in β-DKO mice compared to fl/fl controls (p *=* 0.012), and a loss of structural integrity was also observed in skeletal muscle from the β-DKO mice. Supplementation of β-DKO mice with insulin restored muscle integrity, strength and expression of 13 and 16 of the dysregulated transcripts in and extensor digitorum longus and soleus muscles, respectively.

**Conclusions:** Insulin insufficiency due to β-cell dysfunction perturbs the structure and function of skeletal muscle. These adverse effects are rectified by insulin supplementation.

## Introduction

Skeletal muscle wasting and weakness (sarcopenia) are serious pathological complications of aging and disease[1]. Sarcopenia is a predictor of poor disease outcomes in diabetes[2], cancer[3], chronic heart failure[4] and pneumonia[5] as well as in subjects without underlying disease[6]. A major contributing factor to muscle atrophy is a reduction in circulating levels of the anabolic hormone insulin to below the required threshold for maintaining glucose and muscle homeostasis, either due to insulin resistance or insufficient insulin signalling[7, 8].

Beyond the use of exogenous insulin, treatments for muscle atrophy in diabetes are limited to diet, exercise and the use of insulin sensitising medications such as metformin. However, these options only partially mitigate muscle loss[9]. Whilst impaired muscle metabolism is acknowledged as a serious problem in people with diabetes, questions remain as to the underlying molecular mechanisms that are responsible for this dysfunction and associated structural changes. Moreover, it remains unclear if skeletal muscle dysfunction is a cause or consequence of insulin resistance and disease progression.

Skeletal muscle, which is the main site of glucose disposal, plays a critical role in maintaining glycaemic control[10, 11]. Impaired skeletal muscle metabolism and significant loss of muscle mass due to increased breakdown of myofibrillar proteins, together with reduced exercise capacity, are common in patients with diabetes[12]. Muscle atrophy due to an imbalance in contractile protein synthesis and degradation is also triggered by diabetes[13]. However, there is also evidence that muscle atrophy in diabetes is primarily driven by decreased protein synthesis rather than increased proteolysis[8].

Characterisation of the molecular mechanism(s) by which insufficient insulin action disrupts muscle homeostasis *in vivo* is complicated by limitations in currently available animal models. The most common mouse models of diabetes, the *db/db* mouse and the diet induced obesity (DIO) mouse are primarily models of insulin resistance, with secondary compensatory hyperinsulinemia that exceeds physiological levels found in humans. Chemical destruction of pancreatic β-cells in rats and mice with streptozotocin and muscle-specific deletion of the insulin receptor in mice have provided useful insights, but these are models of a total absence of insulin action. Mice with decreased insulin gene dosage have decreased muscle mass[14] and no change in strength[15], but the transcriptional and functional implications of this genetic deletion are not understood, and it is not known if restoration of circulating insulin levels can reverse this phenotype. Mice with skeletal muscle specific deletion of the insulin receptor[16] have reduced muscle mass and function due to decreased protein synthesis[8] as well as a shift in the disposal of glucose from muscle to adipose tissue[17].

We have previously described a mouse model of pancreatic β-cell dysfunction due to conditional deletion of the cholesterol transporters, ABCA1 and ABCG1 (β-DKO mice)[18]. These mice have approximately half of the circulating insulin levels of littermate fl/fl controls[18]. They also lack the first phase of insulin secretion in response to glucose and have markedly decreased skeletal muscle mass and increased fat mass due to a shift in glucose disposal from muscle to adipose tissue, which can be normalised by supplementation with insulin[18].

This study investigates the consequences of the loss of muscle mass in these mice by asking whether the structure and function of the remaining skeletal muscle is impaired. We investigated the direct effects of insulin insufficiency in the animals by quantifying changes in skeletal muscle gene transcription and morphology, as well as exercise capacity and muscle strength. Evidence that insulin supplementation partly restores skeletal muscle homeostasis in these mice is also presented.

## Materials and Methods

### Animal studies

β-DKO and fl/fl control mice were generated as described previously[18]. All experiments were performed on 12-week-old male mice. The animals were maintained on a standard chow diet in a specific pathogen-free laboratory with a 12 h light/dark cycle. All animal experiments were approved by the Animal Ethics Committee, UNSW Sydney, Ethics Number 16/167B.

### Insulin supplementation

β-DKO mice were supplemented with insulin or PBS (control) in mini-osmotic pumps (Model 1004, Alzet, Cupertino, CA) as described previously[18]. Fl/fl littermate control mice received pumps containing PBS. The pumps were loaded with Humulin R (0.1 U/day, Eli Lily, Indianapolis, IN) or an equivalent volume of PBS prior to subcutaneous implantation between the shoulder blades of anesthetized mice. The mice were euthanized by cervical dislocation 28 days post-pump implantation.

### Tissue collection

The gastrocnemius, soleus, and extensor digitorum longus (EDL) muscles were rapidly excised after euthanasia. Muscles were either used fresh for biomechanics experiments, fixed overnight at 4 °C for microscopy and histology experiments or snap-frozen in liquid nitrogen and stored at −80 °C until further use.

### Gene expression studies

Total RNA was extracted from soleus and EDL muscle using an miRNeasy mini kit (Qiagen, Venlo, Netherlands) according to the manufacturer’s protocol. To ensure RNA was of sufficient quantity and quality for analysis EDL samples from three separate animals were pooled and the experiment performed in duplicate and soleus samples from two separate animals were pooled and the experiment performed in triplicate. RNA was analysed for quality and integrity and sequenced using Agilent 2100 Bioanalyzer (Agilent Technologies, Santa Clara, CA). Libraries were prepared from 500 ng of total RNA/sample using TruSeq Stranded mRNA-seq prep. Each library was single-read sequenced (1×75 bp) using NextSeq 500 High Output flow cell (60 million reads per sample). RNA-seq datasets have been deposited into ENA (https://www.ebi.ac.uk/ena/browser/) with a primary accession number E-MTAB-13315.

### Bioinformatic analysis

RNA-seq reads were first assessed for quality using the FastQC tool (v0.11.8) (www.bioinformatics.babraham.ac.uk/projects/fastqc/). The tool Salmon was used for quantifying transcript abundance from RNA-seq reads[19]. A mouse transcriptome which contained the super-set of all transcripts coding sequences resulting from Ensembl gene predictions (Mus musculus.GRCm38) was downloaded and converted into a Salmon index file (ftp://ftp.ensembl.org/pub/release-101/fasta/mus_musculus/cds/). This index file was used as a reference file for the mapping and quantification step. Transcript level quantifications were converted to gene level quantifications for downstream differential expression analysis.

Differentially expressed genes across all of the conditions were identified using the Bioconductor package edgeR v3.28.0 in the R programming environment[20]. Read counts were normalised using the between lane normalisation of EDASeq with upper quartile normalisation[21]. The R package, RUV, of Risso et al. was used to remove unwanted variation caused by batch effects using empirical control genes[22]. Differentially expressed genes identified with a false discovery rate (FDR) < 0.1 cut-off were considered to be significantly upregulated or downregulated. Transcripts where a replicate had a read value of 0 were excluded from analysis. The Mouse Portal within the Rat Genome Database[23] was used to identify canonical pathways associated with the differentially expressed genes.

### EchoMRI

The percentage of body fat and lean tissue mass of the animals was determined using an EchoMRI-900 Body Composition Analyzer (Biological Resources Imaging Laboratory, UNSW Sydney). Body fat percentage was calculated as:

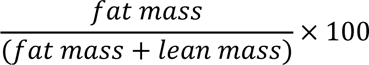

and lean mass percentage as:

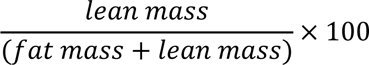

Body composition measurements were performed at baseline (one day prior to insulin supplementation) and after 28 days.

### Transmission electron microscopy (TEM)

For TEM, the EDL was fixed in a 2% (w/v) glutaraldehyde solution with 0.1 M sodium cacodylate, pH 7.4. The samples were then fixed in 1% (w/v) osmium tetroxide (OsO_4_) for 1 h and dehydrated sequentially in acetone (25%, 50%, 75% and 100% (v/v)) twice over 24 h. The EDL was then embedded in 100% resin at 60 °C for 24 h. Ultrathin sections (70 nm) were cut, stained with 2% (v/v) aqueous uranyl acetate and lead citrate and examined on a JEOL TEM-1400 transmission electron microscope (Electron Microscope Unit, Mark Wainwright Analytical Centre, UNSW Sydney, Australia). Electron micrographs were taken at 10,000-100,000× magnification. Semi-quantitative scoring of mitochondrial quality was performed as previously described[24] by a researcher blinded to genotype and treatment, with mitochondria categorized as normal, abnormal or disintegrated (Supplementary Figure 1).

### Exercise protocol

For exercise capacity studies, mice were acclimatised for three consecutive days (20 min/day) on a rodent treadmill (Exer 3/6 Treadmill, Columbus Instruments, Columbus, OH). On the first day, the treadmill speed was set to a warm-up period of running at 5 m/min for 10 min. On the following training days, the initial speed was increased by 1 m/min every min to 15 m/min after the 10 min warm-up period. The mice were then rested for 2 days before the experimental run. On the day of the treadmill studies, mice were run on the treadmill at an initial speed of 15 m/min after the 10 min warm-up period. After 30 min, the speed was increased by 1 m/min every 15 min. Running was confirmed visually, and mice that stopped running were gently pushed back onto the belt with a paper towel. The mice were run to exhaustion, as judged by refusal to remain on the treadmill belt after 5 consecutive failed attempts to return mice to the belt. The assessor was blinded to phenotype. Exercise capacity was calculated as performed work using the following formula:

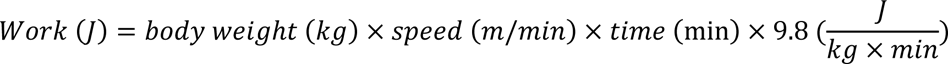

Exercise studies were performed at baseline (2 days prior to insulin supplementation, 1 day prior to EchoMRI) and 28 days after insulin pump implantation.

### Single muscle fibre biomechanical properties

Muscle strength was assessed by contractile performance assessment. Single fibre segments were manually isolated from dissected EDL muscles and myofibrillar Ca^2+^ sensitivity of the contractile apparatus quantified using the *MyoRobot* automated biomechatronics platform as described previously[25, 26]. Five to ten single fibres were analysed for each individual muscle sample. After mild membrane permeabilization with saponin, Ca^2+^-induced force responses were induced by sequentially bathing the skinned fibre in solutions with increasing Ca^2+^ concentrations using defined mixtures of Ca^2+^ free and Ca^2+^-saturated internal solutions[25, 26]. After reconstructing a sigmoidal curve from the steady-state force levels over each pCa, the pCa_50_ was extracted. To control for variability in baseline readings across experiments, values were normalised to maximum force.

### Statistical analyses

Data were assessed for normality using the D’Agostino-Pearson normality test. Differences between the groups were compared using the Wilcoxon test for paired analyses and Mann-Whitney U test for unpaired analyses or one way ANOVA, as appropriate. A value of *p* < 0.05 was considered to be statistically significant. GraphPad Prism software (version 9) was used for statistical analyses.

## Results

### Insulin insufficiency regulates gene expression in skeletal muscle

The impact of insufficient insulin on the relative abundance of skeletal muscle fibre type was assessed in the gastrocnemius of fl/fl and β-DKO mice. β-DKO mice had significantly reduced numbers of Myh1 positive (glycolytic) fibres (72.60±8.56% v 55.20±5.76, p < 0.01; Figure 1A-C) and significantly increased numbers of Mhy7 positive (oxidative) fibres (4.36±1.09 v 8.04±0.98, p < 0.005; Figure 1D-F) compared to fl/fl controls.

**Figure 1:**
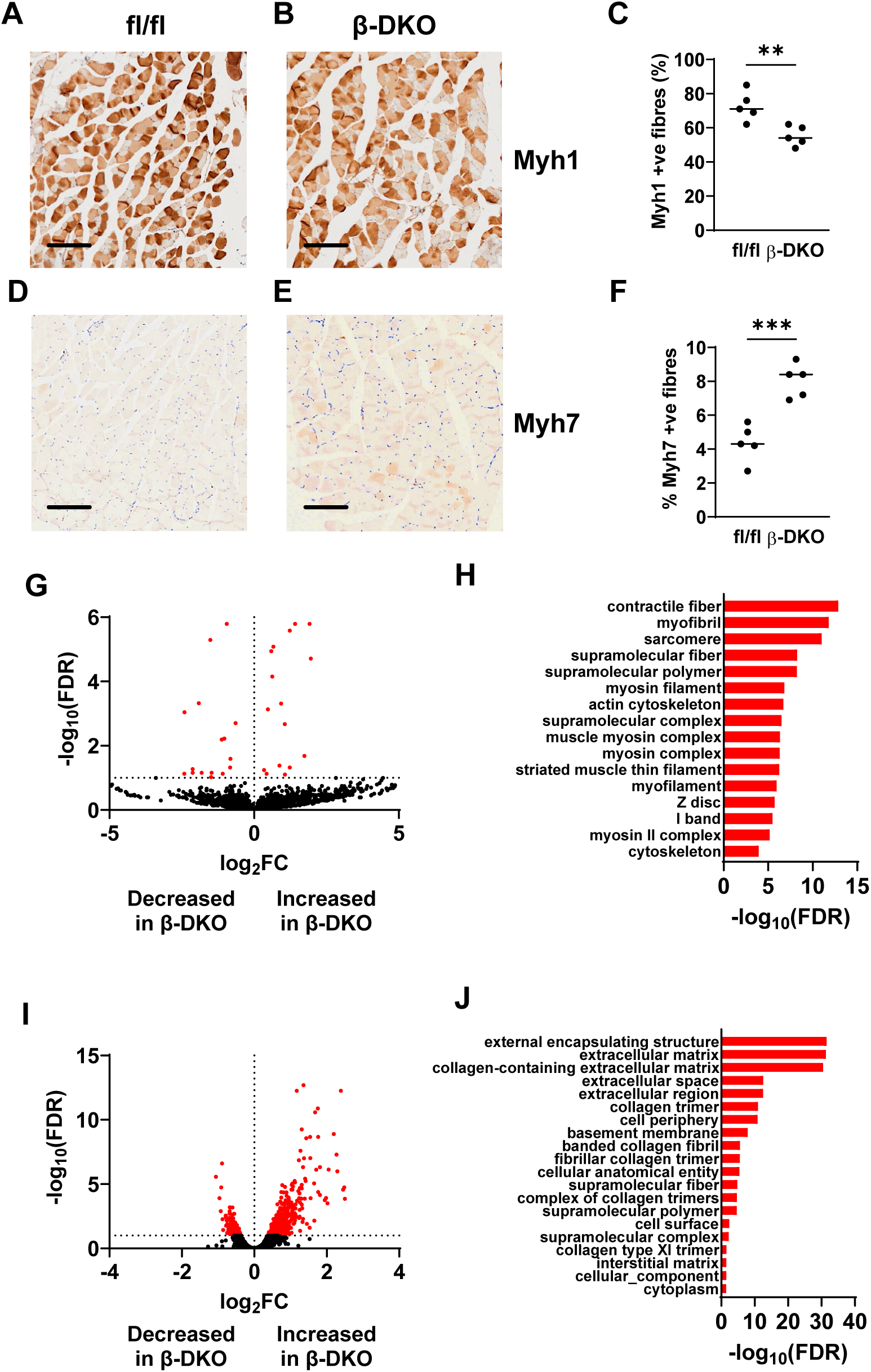
Insulin insufficiency alters expression of transcripts involved in skeletal muscle structure and function. Gastrocnemius muscle was isolated from fl/fl control and β-DKO mice and stained for either (**A-C**) Myh1 or (**D-F**) Myh7. Bulk RNA sequencing was conducted on isolated (**G**) EDL and (**I**) soleus muscles from control fl/fl and β-DKO mice to identify differentially regulated transcripts. Gene Ontology cellular component analysis for (**H**) EDL and (**J**) soleus datasets was obtained using the Mouse Portal of the Rat Genome Database[23]. ** p < 0.01, *** p < 0.005. Scale bar = 200 µm.

To further examine the molecular consequences of insufficient insulin on skeletal muscle gene expression, we performed RNA sequencing on pooled EDL samples. A total of 32 transcripts were differentially regulated between the fl/fl control mice and β-DKO mice that did not receive insulin supplementation (Figure 1G; Supplementary Table 1). Gene Ontology cellular component analysis identified significant differential expression of various components of the contractile apparatus and cytoskeleton (Figure 1H). Biological process analysis indicated significant alterations in pathways involved in contraction and movement (Supplementary Table 5).

RNA sequencing was also performed on soleus samples. A total of 410 transcripts were differentially regulated between the fl/fl control mice and β-DKO mice that did not receive insulin supplementation (Figure 1I; Supplementary Table 2). Gene Ontology cellular component analysis identified significant differential expression of collagen associated extracellular structures (Figure 1J), with significantly increased Col8a2, Col11a1, Col12a1, Col1a2, Col1a1, Col14a1, Col16a1, Col27a1, Col6a2, Col5a2, Col6a1, Col18a1, Col6a3, Col5a1 and Col3a1 transcripts in β-DKO samples relative to fl/fl controls (Supplementary Table 2). Biological process analysis indicated significant alterations in pathways associated with muscle structure and function (Supplementary Table 6).

We next asked what impact insulin supplementation in β-DKO mice had on the expression of muscle transcripts. We used an insulin dosing regimen that restores plasma insulin levels to that of control mice in the fed state[18] (Supplementary Figure 1). Principle component analysis demonstrated more robust clustering of replicates of EDL samples (Figure 2A) than soleus samples (Figure 2B). In EDL samples, a significant difference in the expression of 25 transcripts between β-DKO mice treated with insulin versus PBS was observed (Figure 2C; Supplementary Table 3), with 13 of these transcripts being restored to levels similar to those observed in fl/fl control mice (Figure 2D), including transcripts critical for muscle function and homeostasis such as myoglobin and myosin subunits 1 (*Myh1*) and 4 (*Myh4*). In soleus samples a significant difference in the expression of 76 transcripts between β-DKO mice treated with insulin versus PBS was observed (Figure 2D; Supplementary Table 4), with 16 of these transcripts being restored to levels similar to those observed in fl/fl control mice (Figure 2F).

**Figure 2:**
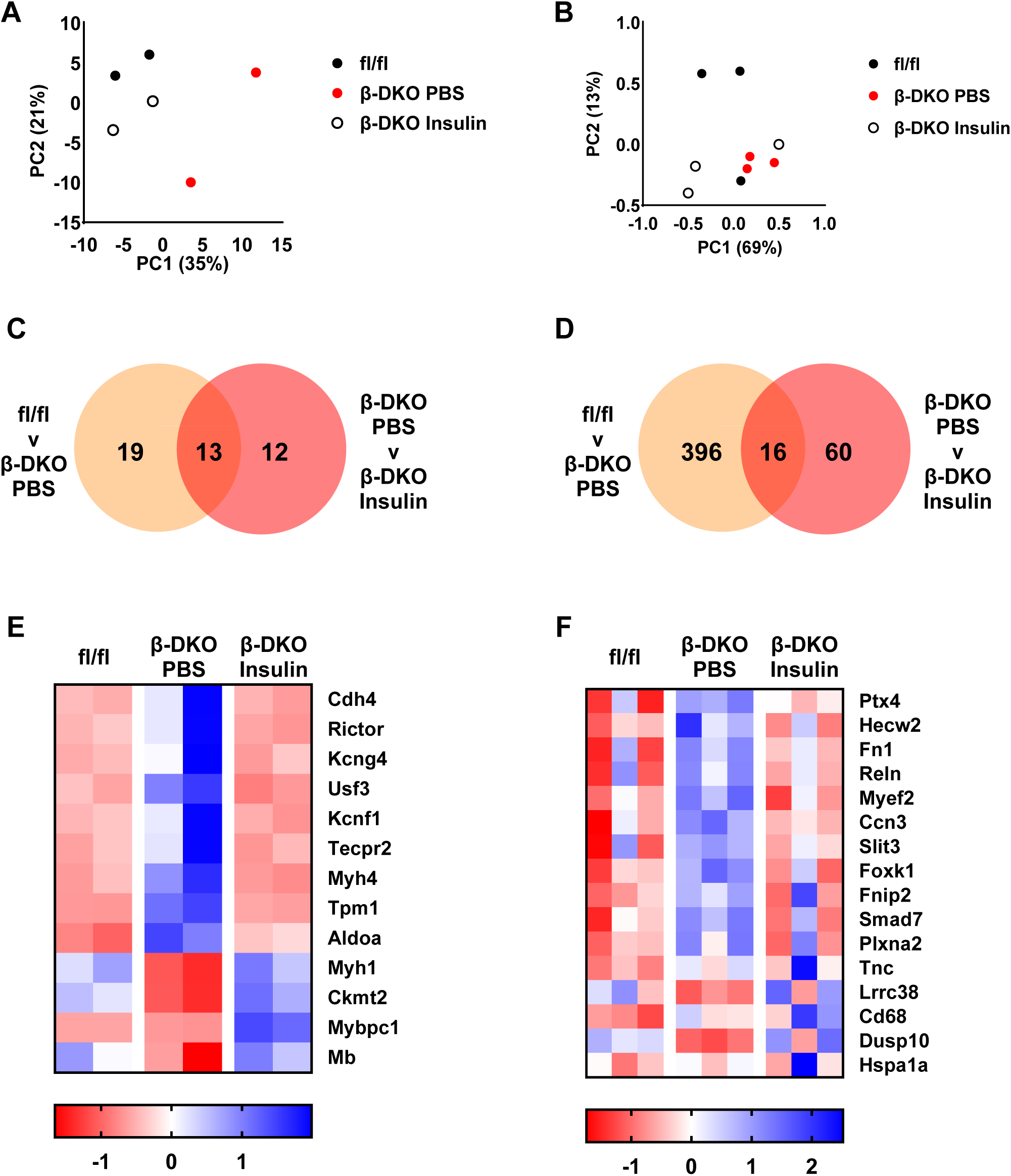
Insulin supplementation normalised the altered expression of transcripts in the EDL and soleus muscles of β-DKO mice. Principal component analysis of sequenced RNA transcripts from (**A**) EDL and (**B**) soleus muscles of fl/fl control (black dots) and β-DKO mice treated with PBS (red dots) or insulin (white dots). Insulin supplementation restored expression of (**C**) 13 transcripts in EDL muscle and (**D**) 16 transcripts in soleus muscle. Heat map analysis of transcripts normalised in (**E**) EDL and (**F**) soleus muscle by insulin treatment.

### Skeletal muscle loss is inhibited by insulin supplementation in β-DKO mice

Similar to our previous findings[18], a comparable increase in body weight over the treatment period was observed for all groups of mice (p < 0.001 for all; Figure 3<u>, B</u>). Lean body mass percentage significantly decreased over the pump implantation period in β-DKO mice treated with PBS (p < 0.005; Figure 3C, D), there was no significant change in lean body mass percentage in β-DKO mice treated with insulin (Figure 3C, D). There was no significant difference in body fat percentage in all groups across the treatment period (Figure 3E, F).

**Figure 3.**
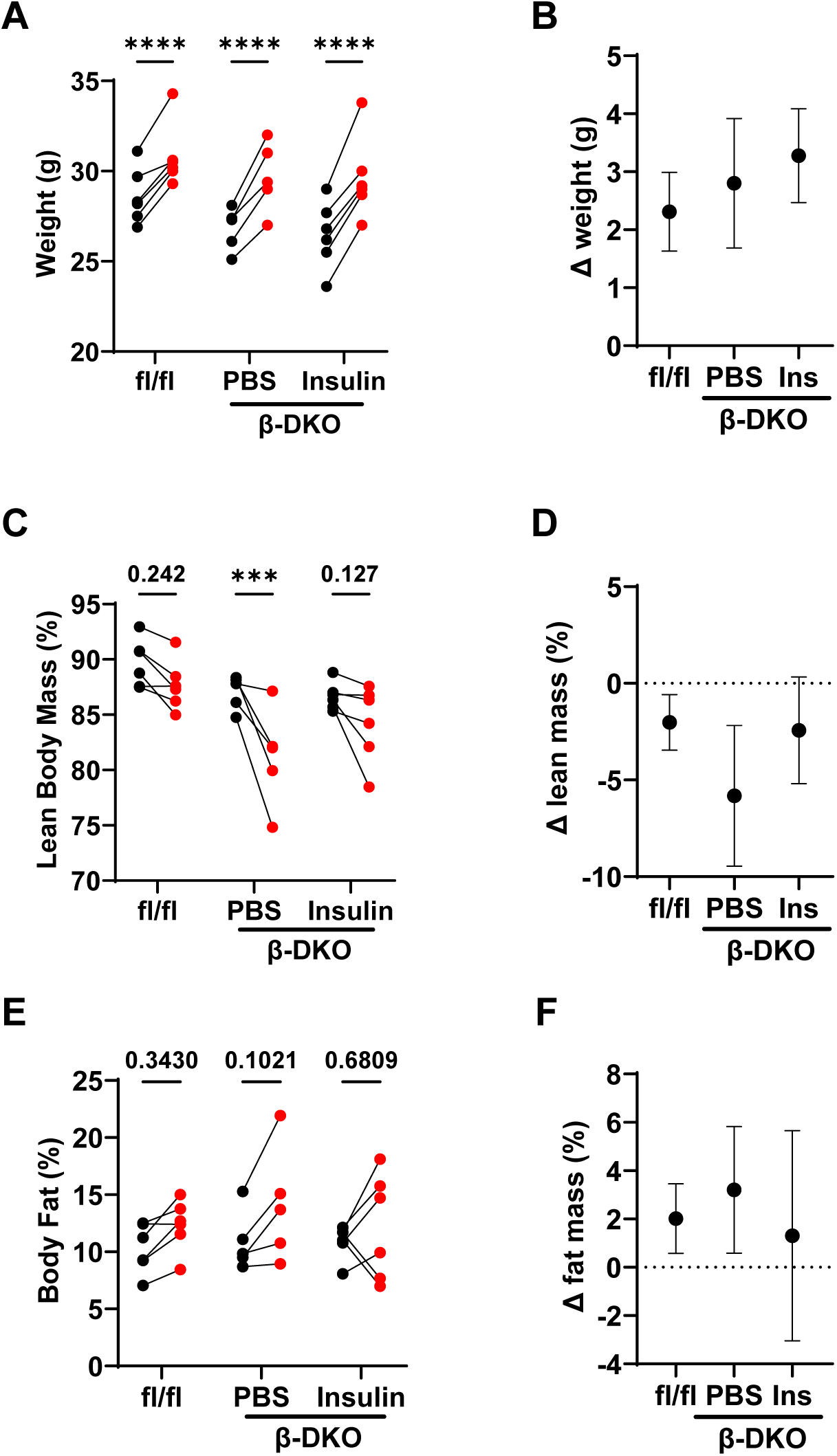
**Insulin supplementation did not alter body weight but** inhibited time-related lean body mass loss. Mice were weighed and subject to echoMRI at baseline (black) and 28 days after pump insertion (red). Change in (**A, B**) body weight, (**C, D**) lean body mass and (**E, F**) body fat in control fl/fl and β-DKO mice treated with PBS or insulin-containing pumps. Data were analysed by two-way ANOVA. Numerical values represent p values. *** p < 0.005, ***** p < 0.001.

### Insulin insufficiency in β-DKO mice drives ultrastructural changes in muscle integrity

Analyses of EDL muscle from control mice by TEM displayed an intact structure with a parallel register of myofibrils and a series of clearly organized sarcomeres with clear Z-lines (Figure 4A). In contrast, muscle from PBS-treated β-DKO mice had severely altered ultrastructure characterized by disorganized myofibrillar register and misaligned Z-lines (Figure 4B). Further, vesicle-like structures were apparent along the Z-lines that protruded into the surrounding myofibrils (Figure 4B, black arrows). Significant restoration of normal ultrastructure was observed in β-DKO mice treated with insulin, with alignment of Z-lines and parallel myofibrillar organisation (Figure 4C).

**Figure 4.**
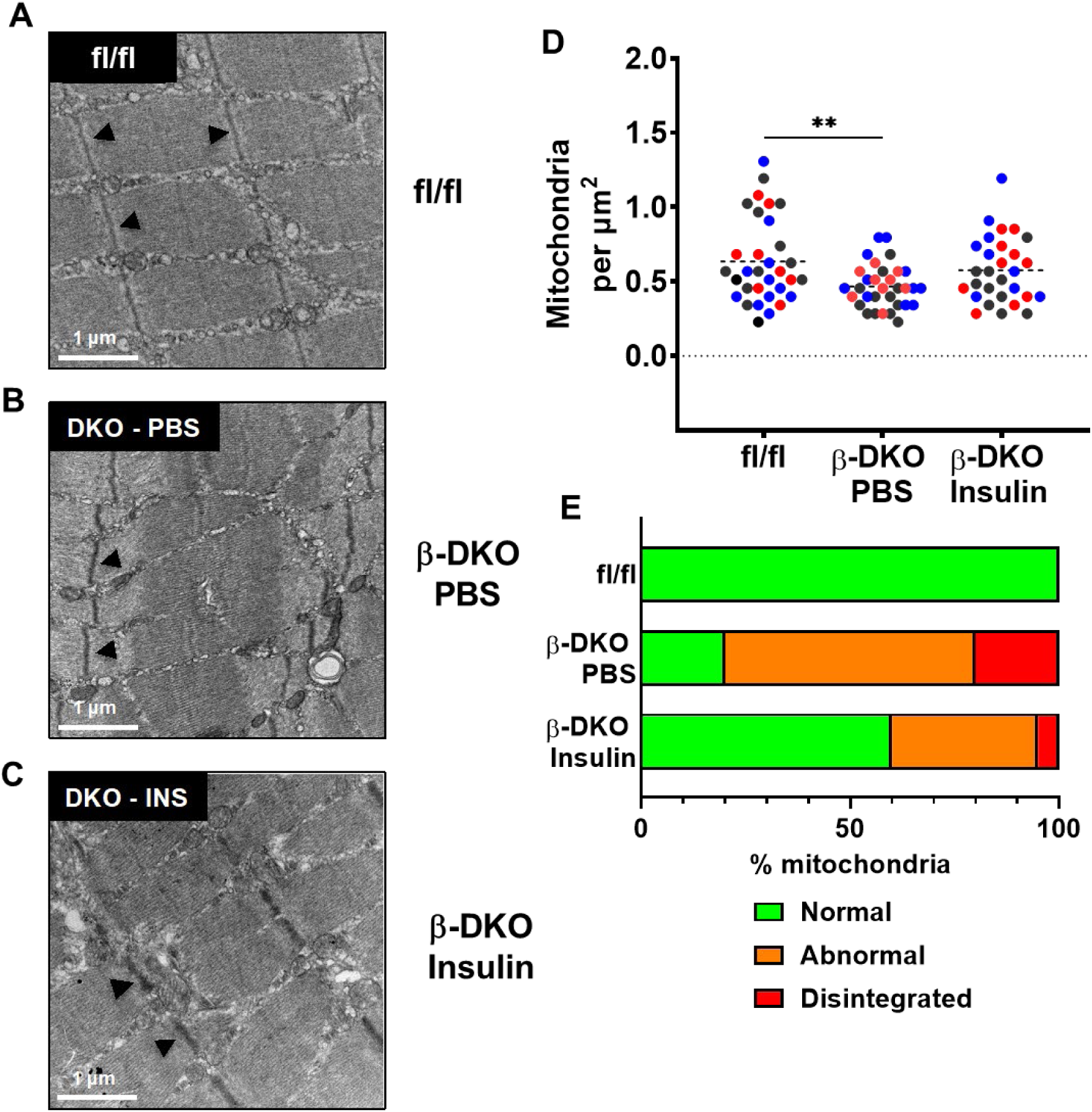
Insulin supplementation restores muscle ultrastructure and mitochondrial morphology. Representative transmission electron microscopy (TEM) images of EDL muscle sections from (**A**) fl/fl control and β-DKO mice treated with (**B**) PBS or (**C**) insulin. Fibre registry and Z-line alignment (black arrowheads) were notably disrupted in the β-DKO PBS mice, and partially restored by insulin treatment. Quantification of mitochondrial (**D**) number and (**E**) quality obtained from multiple fields of view (n = 7-10) for 3 animals per group (each colour represents an independent animal). Dashed lines represent means. **p<0.01. Scale bars = 1 μm.

### Insulin insufficiency decreases mitochondrial number and integrity

Muscle mitochondrial number was significantly decreased in PBS-treated β-DKO mice relative to fl/fl control mice (0.47±0.15 vs 0.63±0.28 per μm^2^, respectively, p<0.01; Figure 4D). Whilst supplementation of β-DKO mice with insulin increased muscle mitochondrial content (0.56±0.22 per μm^2^), this did not reach statistical significance (Figure 4D). To better understand the impact of insulin supplementation on mitochondria, mitochondria were characterised as ‘Normal’, ‘Abnormal’ or ‘Disintegrated’ based on shape and membrane and cristae integrity (Supplementary Figure 2). In muscles isolated from the fl/fl mice, 100% of mitochondria appeared normal (Figure 4E). The proportion of normal mitochondria was significantly reduced in PBS treated β-DKO mice (p<0.001), with most mitochondria classified as abnormal (60%) or disintegrated (20%). Insulin supplementation significantly improved mitochondrial quality (60% normal, 35% abnormal, 5% disintegrated, p<0.001).

### Insulin supplementation restores *in vivo* exercise capacity and single fibre myofibrillar Ca^2+^-sensitivity in β-DKO mice

Prior to pump implantation, low intensity endurance running on a rodent treadmill revealed that exercise capacity was significantly lower in β-DKO mice compared to fl/fl controls (211.8±84.47 vs 366.9±110.7 J for β-DKO and fl/fl mice, respectively; p=0.005; Figure 5A). Significant time-related declines in exercise capacity were observed for both the fl/fl (366.9±110.7 vs 314.5±106.2 J, p<0.01) and PBS treated β-DKO (239.7±113.2 vs 183.7±94.7 J, p<0.005) groups 28 days post pump implantation. Insulin supplementation not only prevented the decline in running performance but significantly increased exercise capacity in β-DKO mice (187.3±42.7 vs 254.9±39.4 J, p<0.005).

**Figure 5.**
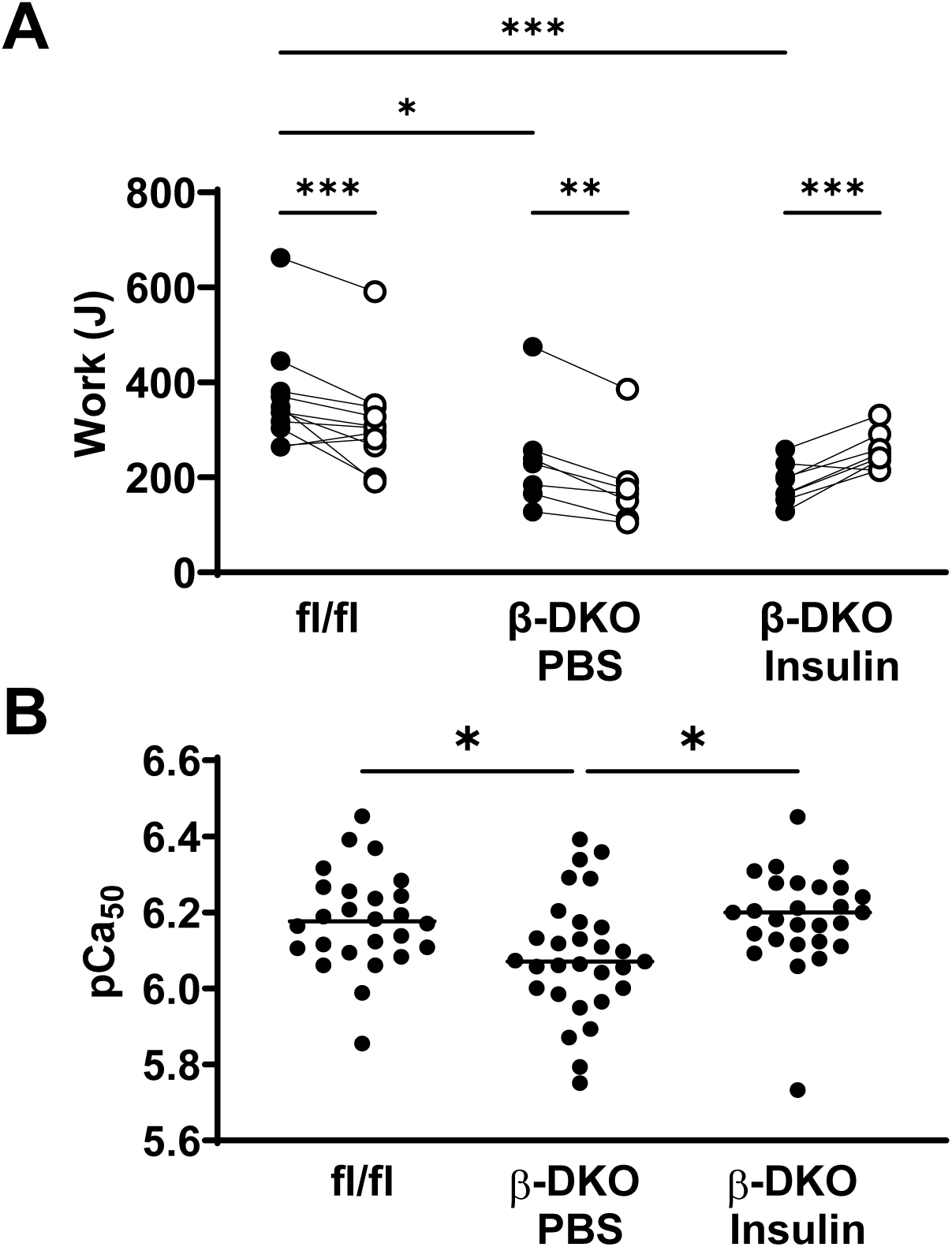
Insulin treatment improves exercise capacity and muscle strength in β-DKO mice. **(A)** Individual running performance was assessed at baseline (black circles) and 28 days post (white circles) pump implantation in fl/fl control and β-DKO mice treated with PBS or insulin. (**B**) Ca^2+^ sensitivity of the contractile apparatus was assessed by subjecting single fibres from the extensor digitorum longus muscle to Ca^2+^-graded force measurements and determining pCa_50_ values. Bars represent mean value. *p<0.05; **p<0.01, ***p<0.001.

In parallel, performance differences were also observed in the Ca^2+^-sensitivity of the contractile apparatus at the single fibre level (Figure 5B). Graded Ca^2+^-induced force measurements performed in isolated EDL skinned fibres demonstrated significant calcium desensitisation in PBS treated β-DKO fibres compared to those from fl/fl control mice (pCa50 = 6.08±0.16 vs 6.18±0.13, respectively; p<0.05). Insulin supplementation corrected the calcium desensitization in β-DKO mice back to control levels (pCa_50_ = 6.19±0.13, p<0.05). There was no change in the maximal force production between the PBS and insulin treated β-DKO mice.

## Discussion

A failure to maintain healthy muscle is a common clinical feature of diabetic skeletal muscle disease and myopathy in type 1 and type 2 diabetes. This condition is characterized by loss of muscle mass, weakness, and overall reduced physical capacity. Distinct changes in skeletal muscle ultrastructure, such as disruption of the fibrillar registry and morphological abnormalities in mitochondria have also been reported in patients with type 1 diabetes[11]. Moreover, accumulation of intracellular lipids has been linked to several pathological complications in skeletal muscle including alterations in insulin sensitivity and signalling. In turn, impaired skeletal muscle metabolism can drive the pathogenesis of insulin resistance and glucose intolerance in diabetes although the underlying mechanisms remain unclear[27]. As the underlying mechanisms responsible for the diminished muscle mass and performance in diabetes are not known, identification of pathophysiological mechanisms and regulation of key metabolic pathways in diabetic muscle will substantially advance current knowledge of the relationship between muscle function and diabetes. This study investigated the effect of β-cell dysfunction and insulin insufficiency on skeletal muscle performance, morphology, mitochondrial structure, and gene expression in mice.

In our previous characterisation of the β-DKO mice, we proposed a shift in metabolic substrates in gastrocnemius muscle due to insufficient circulating insulin levels[18]. This was confirmed by an observed shift from glycolytic to oxidative fibre types in both gastrocnemius and EDL muscles observed in this study. In EDL muscle, which is primarily glycolytic, the significant reduction in circulating insulin levels in the β-DKO mice relative to fl/fl controls resulted in a decrease in the number of myosin heavy chain IIx Myh1 transcript associated with glycolytic fibres which are the predominant constituents of EDL muscle in mice[28, 29]. Supplementation of β-DKO with insulin led to a significant normalisation of the expression of myosin heavy chain isoforms Myh1 and Myh4 as well as myoglobin which plays a critical role in oxygen metabolism in muscle. Whilst Myh2 levels were decreased in the EDL of β-DKO mice relative to fl/fl control, insulin treatment did not reverse this. Insulin has previously been shown to acutely upregulate expression of IIx-associated myosin heavy chain proteins in human vastus lateralis muscle[30]. In contrast, no significant changes in myosin heavy chain transcripts in soleus muscle were observed between groups, presumably as the soleus muscle is already primarily oxidative in nature [31].

The expression of transcripts encoding for several collagen subunits were found to be significantly upregulated in the soleus muscle of β-DKO mice compared to controls. Increased collagen content has described as a feature of insulin resistant skeletal muscle[32], with collagen crosslinking correlated with muscle stiffness [33]. Contrastingly, there were no alterations in the expression of any collagen transcripts in EDL muscle. This finding, together with the myosin findings discussed above, lends weight to the concept that insulin regulation of gene expression differs between muscle types. Previous work has demonstrated that insulin treatment activates distinct signalling events in human type I and type II fibres[34]. Interestingly, whilst insulin treatment improved exercise capacity, it did not reverse the upregulation of collagen transcripts in the soleus muscle of β-DKO mice. This is unlikely to be due to slow turn over collagen in muscle as this study examined expression at the transcript rather than protein level, but rather that either the length of insulin treatment use in this study was too short or that once established, dysregulated collagen expression in skeletal muscle is not reversible. Either way, there are clear implications for clinical outcomes in diabetes and future studies should determine if this process is reversible or not and if it is, the optimal time for intervention to occur.

Insulin treatment resulted in a significant increase in heat shock protein 1A (Hsp70) in soleus muscle. Whilst it is well established that insulin induces expression of Hsp70[35], Hsp70 also regulates skeletal muscle insulin sensitivity [36] and is an important biomarker in contraction injury prevention[37]. As such, modulation of Hsp70 expression has attracted attention as a therapeutic target. Further, Hsp70 regulates FoxO signalling[38], which plays important roles in maintaining muscle energy homeostasis through the control of glycolytic and lipolytic flux, and mitochondrial metabolism[39], as well as regulating diabetes-related muscle atrophy[40]. Insulin treatment also decreased the elevated expression of Foxk1 and Smad7, both of which inhibit the Wnt/β-catenin signalling pathway[41, 42], which plays a critical role in driving myogenesis and inhibiting sarcopenia[43].

Perturbation of skeletal muscle homeostasis in the β-DKO model was associated with decreased exercise capacity and alterations of muscle ultrastructure. The reduced exercise capacity in β-DKO mice and restoration in insulin-treated animals was reflected on the single fibre level with a desensitisation of the contractile apparatus in β-DKO mice compared with controls and restoration following insulin treatment. Previous detailed analyses applying quantitative morphometry to single fibre ultrastructure and simultaneous force recordings in a dystrophic mouse model showed that aberrant myofibrillar alignment is directly correlated with weakness in a graded manner[44]. Whilst insulin supplementation in β-DKO mice resulted in a restoration in muscle organisation, this cannot be directly explained by the transcriptomic data. Rather, it is likely that the action of insulin in this context occurs at a post-translational level, as has been previously characterised in cardiac myocytes[45]. Similarly, insulin signalling has been shown to regulate mitochondrial fusion[46] which plays a critical role in maintaining mitochondrial homeostasis[47].

This study has implications for populations in which β-cell dysfunction can occur prior to insulin resistance[48], such as in patients with type 1 diabetes where loss of skeletal muscle function is more akin to accelerated aging[49]. It suggests that early therapeutic intervention may be the preferred course of treatment for these patients. Indeed, lower BMI and metformin have both been shown to lower the risk of sarcopenia in T2DM[50].

## Acknowledgements

DAAD – Universities Australia Co-operation Scheme #57443180 to OF and PP. The authors acknowledge the facilities and the scientific and technical assistance of the Electron Microscope Unit within the Mark Wainwright Analytical Centre at UNSW Sydney, and in particular the assistance of Mrs Natasha Kapoor-Kaushik.

