## Supplementary Materials for "Effect of insulin insufficiency on ultrastructure and function in skeletal muscle"

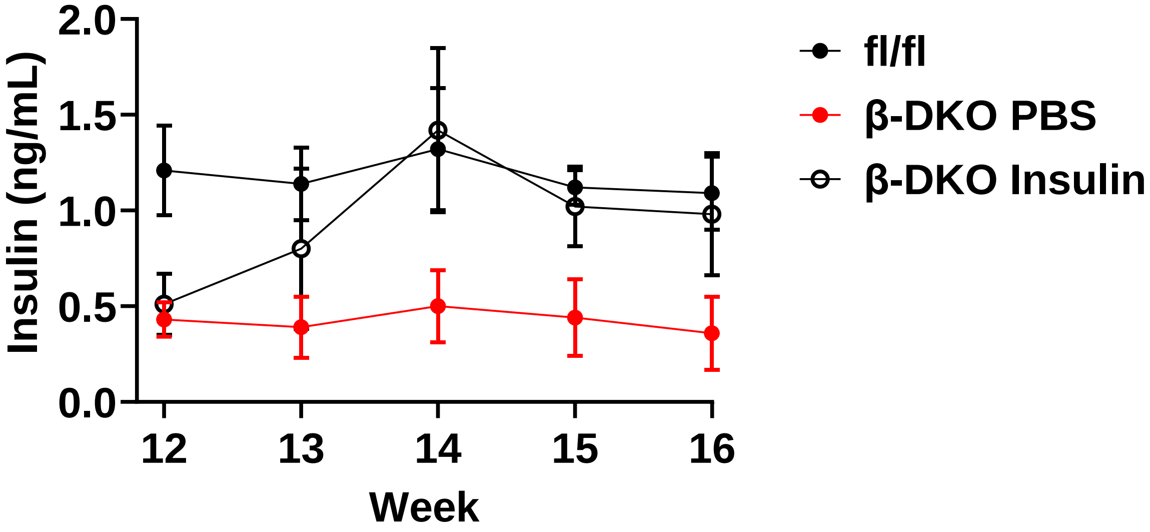


**Supplementary Figure 1: Fed plasma insulin levels in control fl/fl mice and PBS- and insulin-supplemented β-DKO mice.**

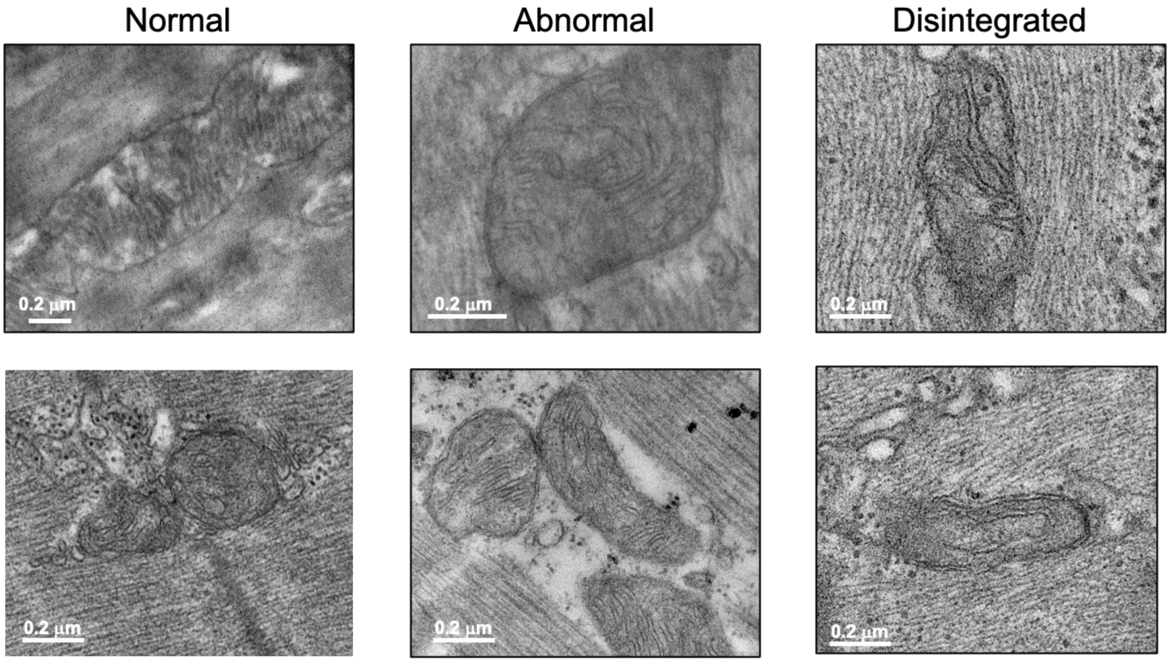


**Supplementary Figure 2: Examples of mitochondria categories used for quantitative analysis of mitochondrial quality in EDL muscle by TEM.**

**Supplementary Table 1: Transcripts differentially expressed in EDL muscle between control fl/fl mice and β-DKO mice.**

| | Fold change | False discovery rate |
|---|---|---|
| **Expression increased in β-DKO vs fl/fl control** | | |
| **cadherin 4\|Cdh4** | 3.892833 | 1.97E-05 |
| **RPTOR independent companion of MTOR, complex 2\|Rictor** | 3.771738 | 1.63E-06 |
| **mab-21-like 1 (C. elegans)\|Mab21l1** | 3.333985 | 0.020934 |
| **potassium voltage-gated channel, subfamily G, member 4\|Kcng4** | 2.663953 | 1.63E-06 |
| **upstream transcription factor family member 3\|Usf3** | 2.35029 | 2.64E-06 |
| **potassium voltage-gated channel, subfamily F, member 1\|Kcnf1** | 2.348649 | 0.048006 |
| **S-adenosylmethionine decarboxylase 1\|Amd1** | 2.086871 | 0.079413 |
| **tectonin beta-propeller repeat containing 2\|Tecpr2** | 2.083586 | 0.002116 |
| **high mobility group AT-hook 2\|Hmga2** | 1.902717 | 0.000487 |
| **serine/threonine kinase 11\|Stk11** | 1.83706 | 0.041278 |
| **myosin, heavy polypeptide 4, skeletal muscle\|Myh4** | 1.582213 | 8.36E-06 |
| **actinin alpha 3\|Actn3** | 1.539407 | 7.1E-05 |
| **tropomyosin 1, alpha\|Tpm1** | 1.507017 | 1.14E-05 |
| **creatine kinase, muscle\|Ckm** | 1.38831 | 0.000742 |
| **muscle glycogen phosphorylase\|Pygm** | 1.349962 | 0.074726 |
| **aldolase A, fructose-bisphosphate\|Aldoa** | 1.267484 | 0.058175 |
| **Expression decreased in β-DKO vs fl/fl control** | | |
| **myosin, heavy polypeptide 1, skeletal muscle, adult\|Myh1** | 1.559699 | 0.001992 |
| **glutathione synthetase\|Gss** | 1.762453 | 0.02551 |
| **creatine kinase, mitochondrial 2\|Ckmt2** | 1.776419 | 0.048006 |
| **myosin binding protein C, slow-type\|Mybpc1** | 1.927011 | 1.63E-06 |
| **actinin alpha 2\|Actn2** | 2.042271 | 0.006034 |
| **protein tyrosine phosphatase, receptor type, Q\|Ptprq** | 2.124977 | 0.074726 |
| **myoglobin\|Mb** | 2.171065 | 0.006388 |
| **EPS8-like 2\|Eps8l2** | 2.766763 | 0.068619 |
| **E74-like factor 1\|Elf1** | 2.816357 | 0.094722 |
| **myosin, heavy polypeptide 2, skeletal muscle, adult\|Myh2** | 2.851151 | 5.18E-06 |
| **spectrin repeat containing, nuclear envelope 1\|Syne1** | 3.512969 | 0.068619 |
| **predicted gene 11127\|Gm11127** | 3.762789 | 0.000473 |
| **leukocyte receptor cluster (LRC) member 8\|Leng8** | 4.360866 | 0.068619 |
| **lactate dehydrogenase B\|Ldhb** | 4.362688 | 0.053787 |
| **ankyrin repeat and zinc finger domain containing 1\|Ankzf1** | 5.299245 | 0.000905 |
| **Ras association (RalGDS/AF-6) domain family (N-terminal) member 8\|Rassf8** | 5.336577 | 0.074726 |

**Supplementary Table 2: Transcripts differentially expressed in soleus muscle between control fl/fl mice and β-DKO mice.**

| | Fold change | False discovery rate |
|---|---|---|
| **Expression increased in β-DKO vs fl/fl control** | | |
| **metallothionein 2\|Mt2** | 315.887858 | 0.00014109 |
| **predicted gene 45799\|Timm10b** | 303.402064 | 1.8111E-05 |
| **tissue inhibitor of metalloproteinase 1\|Timp1** | 281.767758 | 2.6728E-05 |
| **nuclear receptor subfamily 4, group A, member 3\|Nr4a3** | 244.241966 | 5.5621E-13 |
| **collagen, type VIII, alpha 2\|Col8a2** | 195.402066 | 1.013E-06 |
| **fibromodulin\|Fmod** | 186.041089 | 5.054E-08 |
| **tenascin C\|Tnc** | 156.304396 | 1.2894E-09 |
| **collagen, type XI, alpha 1\|Col11a1** | 112.130395 | 7.4789E-07 |
| **angiopoietin-like 7\|Angptl7** | 105.005396 | 0.00031537 |
| **integrin alpha M\|Itgam** | 95.5273219 | 0.00012207 |
| **thrombospondin 1\|Thbs1** | 87.7744799 | 0.00017669 |
| **chondroadherin\|Chad** | 64.5891843 | 6.8925E-05 |
| **collagen, type XII, alpha 1\|Col12a1** | 64.5152457 | 4.822E-07 |
| **pentraxin 4\|Ptx4** | 58.0948417 | 2.155E-09 |
| **chemokine (C-C motif) ligand 6\|Ccl6** | 57.7502693 | 4.2002E-05 |
| **hedgehog interacting protein-like 1\|Hhipl1** | 56.9293577 | 1.3043E-11 |
| **cellular communication network factor 2\|Ccn2** | 51.1747797 | 7.6078E-07 |
| **erythroferrone\|Erfe** | 47.8343156 | 2.6255E-11 |
| **FBJ osteosarcoma oncogene\|Fos** | 45.0717692 | 0.00688853 |
| **tenomodulin\|Tnmd** | 44.9298892 | 8.8498E-06 |
| **sodium channel, voltage-gated, type V, alpha\|Scn5a** | 35.7383512 | 0.00012335 |
| **zinc finger protein 697\|Zfp697** | 35.1236675 | 9.4497E-08 |
| **versican\|Vcan** | 34.5544107 | 2.155E-09 |
| **cartilage intermediate layer protein 2\|Cilp2** | 33.7605039 | 0.02442613 |
| **cartilage oligomeric matrix protein\|Comp** | 28.3777672 | 0.01256866 |
| **lymphatic vessel endothelial hyaluronan receptor 1\|Lyve1** | 27.2160277 | 2.7314E-09 |
| **matrix metallopeptidase 19\|Mmp19** | 26.0262394 | 5.6856E-06 |
| **embigin\|Emb** | 25.2770684 | 3.8547E-06 |
| **proteoglycan 4 (megakaryocyte stimulating factor, articular superficial zone protein)\|Prg4** | 24.84306 | 0.00013438 |
| **runt related transcription factor 1\|Runx1** | 23.5937908 | 9.8254E-08 |
| **mitogen-activated protein kinase kinase kinase 6\|Map3k6** | 22.7629257 | 2.0134E-13 |
| **serine (or cysteine) peptidase inhibitor, clade A, member 3N\|Serpina3n** | 22.0876518 | 4.822E-07 |
| **CD68 antigen\|Cd68** | 22.0040844 | 0.00510463 |
| **early growth response 1\|Egr1** | 21.4435819 | 0.00873695 |
| **coagulation factor II (thrombin) receptor-like 1\|F2rl1** | 20.9021068 | 0.00075681 |
| **thrombospondin 4\|Thbs4** | 20.3645731 | 1.1223E-05 |

| | | |
|---|---|---|
| **cartilage intermediate layer protein, nucleotide pyrophosphohydrolase\|Cilp** | 20.2733309 | 5.6165E-10 |
| **integrin, beta-like 1\|Itgbl1** | 20.267565 | 0.00039092 |
| **fibronectin 1\|Fn1** | 20.262844 | 2.1655E-05 |
| **salt inducible kinase 1\|Sik1** | 19.9031347 | 0.00013438 |
| **latent transforming growth factor beta binding protein 2\|Ltbp2** | 18.7408172 | 2.5832E-08 |
| **insulin receptor substrate 2\|Irs2** | 18.4489986 | 0.0431585 |
| **glutamine fructose-6-phosphate transaminase 2\|Gfpt2** | 18.2581954 | 0.00027226 |
| **peroxisome proliferative activated receptor, gamma, coactivator 1 alpha\|Ppargc1a** | 17.9666502 | 2.0132E-05 |
| **CCAAT/enhancer binding protein (C/EBP), delta\|Cebpd** | 17.8678223 | 0.00337059 |
| **collagen, type I, alpha 2\|Col1a2** | 17.7717894 | 0.00100731 |
| **adhesion G protein-coupled receptor D1\|Adgrd1** | 17.6359685 | 5.2755E-05 |
| **epithelial membrane protein 1\|Emp1** | 17.2024533 | 0.00012572 |
| **integrin beta 2\|Itgb2** | 17.0479969 | 0.00425523 |
| **cation channel, sperm associated 4\|Catsper4** | 16.9025847 | 1.4063E-07 |
| **zinc finger protein 469\|Zfp469** | 16.5022511 | 3.4609E-05 |
| **folliculin interacting protein 2\|Fnip2** | 15.1485024 | 0.00019605 |
| **protein phosphatase 1, regulatory subunit 10\|Ppp1r10** | 14.8919847 | 5.5621E-13 |
| **fibrinogen-like protein 2\|Fgl2** | 14.8377892 | 0.00361599 |
| **N-myc downstream regulated gene 4\|Ndrg4** | 13.7856373 | 0.00037263 |
| **CD44 antigen\|Cd44** | 13.6675049 | 0.00077693 |
| **C-type lectin domain family 11, member a\|Clec11a** | 13.6543471 | 0.00230666 |
| **dihydropyrimidinase-like 3\|Dpysl3** | 13.2537405 | 0.00651421 |
| **a disintegrin-like and metallopeptidase (reprolysin type) with thrombospondin type 1 motif, 9\|Adamts9** | 13.2042952 | 7.4983E-06 |
| **periostin, osteoblast specific factor\|Postn** | 13.0331531 | 5.7526E-06 |
| **solute carrier family 15 (H+/peptide transporter), member 2\|Slc15a2** | 12.9428167 | 0.00178861 |
| **collagen, type I, alpha 1\|Col1a1** | 12.9026108 | 0.00267751 |
| **metallothionein 1\|Mt1** | 12.7005705 | 0.01083837 |
| **lysyl oxidase-like 2\|Loxl2** | 12.4237081 | 0.00344522 |
| **prostaglandin I2 (prostacyclin) synthase\|-** | 12.0811996 | 0.00072563 |
| **RAB30, member RAS oncogene family\|Rab30** | 12.0121819 | 0.01248442 |
| **Rho GTPase activating protein 30\|Arhgap30** | 11.9758937 | 0.00084773 |
| **adhesion G protein-coupled receptor E1\|Adgre1** | 11.9512669 | 0.00055322 |
| **Iqcj and Schip1 fusion protein\|Iqschfp** | 11.8948686 | 0.00044205 |
| **predicted gene, 49804\|-** | 11.7480552 | 0.00204405 |
| **lysosomal-associated protein transmembrane 5\|Laptm5** | 11.6359713 | 1.5222E-05 |
| **coagulation factor XIII, A1 subunit\|F13a1** | 11.2919924 | 2.6657E-05 |
| **adenylate cyclase 7\|Adcy7** | 11.2915835 | 8.93E-06 |
| **HECT, C2 and WW domain containing E3 ubiquitin protein ligase 2\|Hecw2** | 11.1516857 | 0.00221707 |
| **v-maf musculoaponeurotic fibrosarcoma oncogene family, protein F (avian)\|Maff** | 11.0823997 | 0.00065029 |
| **ectodermal-neural cortex 1\|Enc1** | 11.0569355 | 0.00120168 |
| **lysophosphatidic acid receptor 1\|Lpar1** | 10.9459954 | 0.00039616 |
| **serine (or cysteine) peptidase inhibitor, clade B, member 8\|Serpinb8** | 10.8024305 | 1.1223E-05 |

| | | |
|---|---|---|
| **cathepsin S\|Ctss** | 10.7273549 | 0.00524671 |
| **fibronectin type III domain containing 1\|Fndc1** | 10.6129895 | 2.6955E-05 |
| **proline arginine-rich end leucine-rich repeat\|Prelp** | 10.6054774 | 0.00274157 |
| **secreted phosphoprotein 1\|Spp1** | 10.5271111 | 0.0002661 |
| **lysyl oxidase\|Lox** | 10.4327288 | 0.00618155 |
| **Ras-related associated with diabetes\|Rrad** | 10.218397 | 0.02559645 |
| **OTU domain containing 1\|Otud1** | 10.1063917 | 0.00045752 |
| **solute carrier family 38, member 2\|Slc38a2** | 10.0500152 | 0.00213847 |
| **period circadian clock 2\|Per2** | 9.65615931 | 0.00817409 |
| **interferon activated gene 205\|Ifi205** | 9.64754521 | 0.00024503 |
| **lectin, galactose binding, soluble 3\|Lgals3** | 9.28103767 | 0.00274157 |
| **fibulin 1\|Fbln1** | 9.23244477 | 0.02012522 |
| **NUAK family, SNF1-like kinase, 2\|Nuak2** | 9.20759265 | 0.00222847 |
| **cyclin D1\|Ccnd1** | 9.17645131 | 0.02798475 |
| **sortilin-related VPS10 domain containing receptor 2\|Sorcs2** | 9.10927714 | 0.00834923 |
| **WEE 1 homolog 1 (S. pombe)\|Wee1** | 9.08699837 | 0.00065029 |
| **peptidase domain containing associated with muscle regeneration 1\|Pamr1** | 9.00126956 | 0.01345282 |
| **guanine nucleotide binding protein, alpha stimulating, olfactory type\|Gnal** | 8.93123087 | 0.00014109 |
| **N-acetyltransferase domain containing 1\|Natd1** | 8.760899 | 0.00010963 |
| **colony stimulating factor 1 receptor\|Csf1r** | 8.76029693 | 0.00284005 |
| **scavenger receptor cysteine rich family, 5 domains\|Ssc5d** | 8.72194282 | 0.0257702 |
| **lysyl oxidase-like 3\|Loxl3** | 8.5914209 | 0.00055322 |
| **nuclear receptor subfamily 4, group A, member 1\|Nr4a1** | 8.51090577 | 0.00027591 |
| **drebrin 1\|Dbn1** | 8.45805515 | 0.00066092 |
| **mal, T cell differentiation protein-like\|Mall** | 8.44506523 | 0.00200937 |
| **elastin microfibril interfacer 2\|Emilin2** | 8.41190081 | 0.00030487 |
| **myelocytomatosis oncogene\|Myc** | 8.40416371 | 0.01651234 |
| **SRY (sex determining region Y)-box 9\|Sox9** | 8.40348936 | 0.04473441 |
| **mannose receptor, C type 2\|Mrc2** | 8.33397958 | 0.00158168 |
| **reelin\|Reln** | 8.3091645 | 0.00891459 |
| **naked cuticle 2\|Nkd2** | 8.26469802 | 0.00102932 |
| **G protein-coupled receptor 153\|Gpr153** | 8.20302939 | 0.00348355 |
| **zinc finger protein 185\|Zfp185** | 8.18927737 | 0.0002661 |
| **collagen, type XIV, alpha 1\|Col14a1** | 8.1175145 | 0.00139387 |
| **Fc receptor, IgG, low affinity III\|Fcgr3** | 7.81441262 | 0.00912103 |
| **mesenteric estrogen dependent adipogenesis\|Medag** | 7.77438536 | 8.009E-05 |
| **collagen, type XVI, alpha 1\|Col16a1** | 7.76486037 | 0.01502998 |
| **zinc finger and BTB domain containing 16\|Zbtb16** | 7.67545137 | 4.3981E-05 |
| **tubulin, beta 6 class V\|Tubb6** | 7.64289183 | 0.00344522 |
| **pyridoxal (pyridoxine, vitamin B6) kinase\|Pdxk** | 7.59887828 | 0.00995548 |
| **LPS-induced TN factor\|Litaf** | 7.59058792 | 0.00657922 |
| **carboxypeptidase X 2 (M14 family)\|Cpxm2** | 7.54651399 | 0.00884467 |
| **keratin 80\|Krt80** | 7.54373805 | 0.00149876 |
| **macrophage galactose N-acetyl-galactosamine specific lectin 2\|Mgl2** | 7.53519209 | 0.01985801 |

| | | |
|---|---|---|
| **feline leukemia virus subgroup C cellular receptor 1\|Flvcr1** | 7.5301775 | 0.02199621 |
| **podoplanin\|Pdpn** | 7.48006556 | 0.00183658 |
| **sushi, von Willebrand factor type A, EGF and pentraxin domain containing 1\|Svep1** | 7.47219164 | 0.00550052 |
| **phosphodiesterase 10A\|Pde10a** | 7.3361616 | 0.00018081 |
| **CCAAT/enhancer binding protein (C/EBP), beta\|Cebpb** | 7.21129504 | 0.00274157 |
| **procollagen lysine, 2-oxoglutarate 5-dioxygenase 2\|Plod2** | 7.07009335 | 0.0014675 |
| **matrix metallopeptidase 2\|Mmp2** | 7.06573722 | 0.0037345 |
| **advillin\|Avil** | 7.05866928 | 1.6508E-05 |
| **platelet factor 4\|Pf4** | 7.02928514 | 0.03585484 |
| **protocadherin 7\|Pcdh7** | 6.91046647 | 0.00074448 |
| **phosphoinositide-3-kinase regulatory subunit 5\|Pik3r5** | 6.90491314 | 0.00607362 |
| **mitogen-activated protein kinase 4\|Mapk4** | 6.81230376 | 0.01201429 |
| **AE binding protein 1\|Aebp1** | 6.79609015 | 0.00039659 |
| **epidermal growth factor receptor\|Egfr** | 6.76756021 | 0.00393043 |
| **mannoside acetylglucosaminyltransferase 4, isoenzyme A\|Mgat4a** | 6.74957771 | 0.00066955 |
| **Ras and Rab interactor 3\|Rin3** | 6.72265403 | 0.00326217 |
| **dermatan sulfate epimerase\|Dse** | 6.70193524 | 0.00271757 |
| **LRRN4 C-terminal like\|Lrrn4cl** | 6.68675287 | 0.01434866 |
| **discoidin domain receptor family, member 2\|Ddr2** | 6.52784821 | 0.02069978 |
| **solute carrier family 20, member 1\|Slc20a1** | 6.48819816 | 0.00031537 |
| **collagen, type XXVII, alpha 1\|Col27a1** | 6.48620469 | 0.00166425 |
| **follistatin-like 1\|Fstl1** | 6.46539975 | 0.00434387 |
| **mesenchyme homeobox 1\|Meox1** | 6.45591563 | 0.00202072 |
| **fibrillin 1\|Fbn1** | 6.42163214 | 4.4461E-05 |
| **integrin alpha 11\|Itga11** | 6.41694295 | 0.00274157 |
| **Ras association and DIL domains\|Radil** | 6.41093768 | 0.00102932 |
| **plexin A2\|Plxna2** | 6.37656903 | 0.01014739 |
| **proacrosin binding protein\|Acrbp** | 6.35586507 | 0.00107002 |
| **ATPase, class V, type 10A\|Atp10a** | 6.16706082 | 0.006084 |
| **stearoyl-Coenzyme A desaturase 1\|Scd1** | 6.14968502 | 0.01532011 |
| **calcium channel, voltage-dependent, beta subunit associated regulatory protein\|Cbarp** | 6.13632974 | 0.01345282 |
| **serine (or cysteine) peptidase inhibitor, clade F, member 1\|Serpinf1** | 6.04494677 | 0.02234566 |
| **regulatory factor X, 2 (influences HLA class II expression)\|Rfx2** | 6.02231851 | 0.04582482 |
| **DENN/MADD domain containing 2A\|Dennd2a** | 5.990666 | 0.00084706 |
| **predicted gene 11756\|Gm11756** | 5.93200883 | 0.00805211 |
| **growth arrest specific 1\|Gas1** | 5.93058597 | 0.00753885 |
| **elongation factor RNA polymerase II\|Ell** | 5.91322391 | 0.00438061 |
| **interleukin 17D\|Il17d** | 5.91250418 | 0.02559645 |
| **plexin A4\|Plxna4** | 5.87885105 | 1.2001E-05 |
| **v-maf musculoaponeurotic fibrosarcoma oncogene family, protein K (avian)\|Mafk** | 5.86462621 | 0.02026195 |
| **collagen, type VI, alpha 2\|Col6a2** | 5.85196462 | 0.00291368 |
| **low density lipoprotein receptor-related protein 1\|Lrp1** | 5.84744648 | 0.00075191 |
| **tumor necrosis factor receptor superfamily, member 25\|Tnfrsf25** | 5.82106598 | 0.02934215 |

| | | |
|---|---|---|
| scleraxis\|Scx | 5.79195978 | 0.00034609 |
| complement component factor h\|Cfh | 5.77817173 | 0.02529444 |
| TBC1 domain family, member 9\|Tbc1d9 | 5.75724638 | 0.01224856 |
| myeloid-associated differentiation marker\|Myadm | 5.74757747 | 0.01393496 |
| cysteine-serine-rich nuclear protein 1\|Csrnp1 | 5.71080554 | 0.00689744 |
| ENSMUSG00000121578 | 5.70323026 | 0.01155858 |
| stromal cell-derived factor 2-like 1\|Sdf2l1 | 5.64213861 | 0.01372805 |
| Kruppel-like factor 3 (basic)\|Klf3 | 5.63196996 | 0.00510463 |
| proprotein convertase subtilisin/kexin type 5\|Pcsk5 | 5.62031347 | 0.00232909 |
| E74-like factor 4 (ets domain transcription factor)\|Elf4 | 5.57009864 | 0.00425721 |
| tenascin XB\|Tnxb | 5.55854386 | 0.00011146 |
| regulator of calcineurin 1\|Rcan1 | 5.53858966 | 4.1168E-05 |
| AXL receptor tyrosine kinase\|Axl | 5.53691521 | 0.00682128 |
| H1.9 linker histone\|H1f9 | 5.52571745 | 0.00773557 |
| thrombospondin 3\|Thbs3 | 5.50847072 | 0.00077693 |
| carboxylesterase 5A\|Ces5a | 5.4748406 | 0.00750199 |
| CD109 antigen\|Cd109 | 5.43003574 | 0.00230666 |
| Rho GTPase activating protein 26\|Arhgap26 | 5.41330348 | 0.02245268 |
| matrix Gla protein\|Mgp | 5.39936149 | 0.02225316 |
| cerebellar degeneration-related 2\|Cdr2 | 5.39077385 | 0.00193926 |
| tubulin, beta 2B class IIB\|Tubb2b | 5.33587057 | 0.00533211 |
| slit guidance ligand 3\|Slit3 | 5.31656619 | 0.00346577 |
| collagen, type V, alpha 2\|Col5a2 | 5.27074042 | 0.00626291 |
| pre B cell leukemia homeobox 3\|Pbx3 | 5.25387109 | 0.01393496 |
| complement component 1, q subcomponent, C chain\|C1qc | 5.25333539 | 0.02483316 |
| strawberry notch 2\|Sbno2 | 5.24913287 | 0.00258716 |
| nuclear factor, interleukin 3, regulated\|Nfil3 | 5.20506727 | 0.01759268 |
| sorbin and SH3 domain containing 3\|Sorbs3 | 5.20192731 | 0.00022721 |
| unc-13 homolog D\|Unc13d | 5.16992642 | 0.00434387 |
| olfactomedin-like 2B\|Olfml2b | 5.15691555 | 0.01393496 |
| chloride intracellular channel 4 (mitochondrial)\|Clic4 | 5.15489816 | 0.00072048 |
| solute carrier family 7 (cationic amino acid transporter, y+ system), member 1\|Slc7a1 | 5.12964325 | 0.00102932 |
| von Willebrand factor A domain containing 1\|Vwa1 | 5.12267261 | 0.00906574 |
| SH3 domain protein D19\|Sh3d19 | 5.1183914 | 0.00425523 |
| interferon activated gene 204\|Ifi204 | 5.08152601 | 0.00556062 |
| paralemmin A kinase anchor protein\|Pakap | 5.08088233 | 0.04110072 |
| alkaline phosphatase, liver/bone/kidney\|Alpl | 5.03502833 | 0.00337059 |
| a disintegrin and metallopeptidase domain 19 (meltrin beta)\|Adam19 | 5.02576571 | 0.0431585 |
| toll-like receptor 4\|Tlr4 | 5.01236395 | 0.00380425 |
| UDP-Gal:betaGlcNAc beta 1,4-galactosyltransferase, polypeptide 6\|B4galt6 | 5.01164896 | 0.00274157 |
| tumor necrosis factor receptor superfamily, member 12a\|Tnfrsf12a | 4.9800393 | 0.01083044 |
| tissue inhibitor of metalloproteinase 2\|Timp2 | 4.85259315 | 0.00907549 |
| transgelin 2\|Tagln2 | 4.82472343 | 0.02934215 |

| | | |
|---|---|---|
| **anthrax toxin receptor 1\|Antxr1** | 4.80748239 | 0.01336038 |
| **sushi-repeat-containing protein, X-linked 2\|Srpx2** | 4.80078628 | 0.03908268 |
| **acetoacetyl-CoA synthetase\|Aacs** | 4.7839415 | 0.00310573 |
| **coactosin-like 1 (Dictyostelium)\|Cotl1** | 4.74828626 | 0.01807052 |
| **vimentin\|Vim** | 4.73968773 | 0.00254365 |
| **vitrin\|Vit** | 4.71461086 | 0.0388031 |
| **tropomyosin 4\|Tpm4** | 4.71160534 | 0.02049759 |
| **S100 calcium binding protein A10 (calpactin)\|S100a10** | 4.70823788 | 0.00968023 |
| **poliovirus receptor\|Pvr** | 4.70113839 | 0.01004436 |
| **platelet-derived growth factor receptor-like\|Pdgfrl** | 4.683342 | 0.00626291 |
| **Fraser extracellular matrix complex subunit 1\|Fras1** | 4.6754029 | 0.00141916 |
| **lecithin cholesterol acyltransferase\|Lcat** | 4.62697522 | 0.03951736 |
| **AHNAK nucleoprotein 2\|Ahnak2** | 4.59316187 | 0.00750199 |
| **ENSMUSG00000095041** | 4.58785789 | 0.00525878 |
| **CDP-diacylglycerol synthase 1\|Cds1** | 4.58400127 | 0.04110072 |
| **annexin A1\|Anxa1** | 4.57150044 | 0.00223869 |
| **cysteine and glycine-rich protein 3\|Csrp3** | 4.55811356 | 0.0280054 |
| **protein phosphatase 1, regulatory subunit 13 like\|Ppp1r13l** | 4.55148548 | 0.01079525 |
| **biglycan\|Bgn** | 4.52591981 | 0.01393496 |
| **death associated protein kinase 1\|Dapk1** | 4.49901288 | 0.00750019 |
| **laminin, gamma 2\|Lamc2** | 4.48323987 | 0.00274157 |
| **zinc finger and BTB domain containing 26\|Zbtb26** | 4.48093318 | 0.03756582 |
| **transforming growth factor, beta induced\|Tgfbi** | 4.47854087 | 0.01336038 |
| **microtubule-associated protein 1B\|Map1b** | 4.4709083 | 0.00393043 |
| **integrin, alpha 10\|Itga10** | 4.46649125 | 0.03982991 |
| **Sec24 related gene family, member D (S. cerevisiae)\|Sec24d** | 4.42938814 | 0.01345282 |
| **collagen, type VI, alpha 1\|Col6a1** | 4.4147565 | 0.0099896 |
| **collagen, type XVIII, alpha 1\|Col18a1** | 4.39078253 | 0.01174358 |
| **NLR family, CARD domain containing 5\|Nlrc5** | 4.37491322 | 0.03588503 |
| **epidermal growth factor-containing fibulin-like extracellular matrix protein 1\|Efemp1** | 4.35614105 | 0.00817409 |
| **phosphodiesterase 5A, cGMP-specific\|Pde5a** | 4.35052855 | 0.00890655 |
| **WD repeat domain 81\|Wdr81** | 4.34261868 | 0.03509434 |
| **microtubule associated monooxygenase, calponin and LIM domain containing -like 1\|Micall1** | 4.33564834 | 0.00036909 |
| **ret proto-oncogene\|Ret** | 4.3256681 | 0.03990217 |
| **receptor-interacting serine-threonine kinase 3\|Ripk3** | 4.31490109 | 0.03677365 |
| **a disintegrin-like and metallopeptidase (reprolysin type) with thrombospondin type 1 motif, 8\|Adamts8** | 4.28204334 | 0.02341984 |
| **carboxylesterase 2C\|Ces2c** | 4.2809602 | 0.03002208 |
| **Jun dimerization protein 2\|Jdp2** | 4.26638065 | 0.01653921 |
| **fibroblast growth factor receptor 3\|Fgfr3** | 4.26278108 | 0.00772354 |
| **collagen, type VI, alpha 3\|Col6a3** | 4.25156346 | 0.01735642 |
| **ring finger protein 144B\|Rnf144b** | 4.25147582 | 0.01079525 |
| **annexin A2\|Anxa2** | 4.23569249 | 0.00434387 |
| **proprotein convertase subtilisin/kexin type 6\|Pcsk6** | 4.21986739 | 0.01133451 |
| **microfibrillar associated protein 5\|Mfap5** | 4.21593664 | 0.00706405 |

| | | |
|---|---|---|
| **endothelin receptor type B\|Ednrb** | 4.20462609 | 0.0431585 |
| **prostate transmembrane protein, androgen induced 1\|Pmepa1** | 4.19813485 | 0.02012522 |
| **RNA binding motif protein 12 B1\|Rbm12b1** | 4.1917968 | 0.02341984 |
| **FYVE, RhoGEF and PH domain containing 3\|Fgd3** | 4.19105663 | 0.00825343 |
| **EYA transcriptional coactivator and phosphatase 1\|Eya1** | 4.1765802 | 0.0162846 |
| **transferrin receptor\|Tfrc** | 4.16271718 | 0.01014739 |
| **chemokine-like receptor 1\|Cmklr1** | 4.15864414 | 0.01226934 |
| **RIKEN cDNA 1110065P20 gene\|1110065P20Rik** | 4.12216788 | 0.03158457 |
| **RAB3A interacting protein (rabin3)-like 1\|Rab3il1** | 4.10582684 | 0.02988793 |
| **zinc finger SWIM-type containing 6\|Zswim6** | 4.09863205 | 0.01130906 |
| **ski sarcoma viral oncogene homolog (avian)\|Ski** | 4.09195668 | 0.02679512 |
| **ADP-ribosylation factor-like 13B\|Arl13b** | 4.09171313 | 0.00753885 |
| **collagen, type V, alpha 1\|Col5a1** | 4.06489257 | 0.0079776 |
| **forkhead box O1\|Foxo1** | 4.0504303 | 0.0306347 |
| **tumor necrosis factor receptor superfamily, member 23\|Tnfrsf23** | 4.03722408 | 0.04764597 |
| **UDP-glucose dehydrogenase\|Ugdh** | 4.0301866 | 0.00264925 |
| **TOG array regulator of axonemal microtubules 2\|Togaram2** | 4.02596612 | 0.01224856 |
| **collagen, type III, alpha 1\|Col3a1** | 4.01251179 | 0.01336038 |
| **pleckstrin homology domain containing, family F (with FYVE domain) member 1\|Plekhf1** | 4.00483924 | 0.03998977 |
| **cellular communication network factor 3\|Ccn3** | 3.99111868 | 0.03542224 |
| **inter-alpha (globulin) inhibitor H5\|Itih5** | 3.95945166 | 0.01248442 |
| **transcriptional and immune response regulator\|Tcim** | 3.95046319 | 0.00689843 |
| **tensin 3\|Tns3** | 3.94656826 | 0.03310089 |
| **calcium channel, voltage-dependent, T type, alpha 1G subunit\|Cacna1g** | 3.94483098 | 0.00682128 |
| **expressed sequence C77080\|C77080** | 3.9437829 | 0.01079525 |
| **signal-regulatory protein alpha\|Sirpa** | 3.92615826 | 0.03644731 |
| **heat shock protein 1A\|Hspa1a** | 3.90723182 | 0.00639162 |
| **forkhead box K1\|Foxk1** | 3.88326081 | 0.01182355 |
| **sperm associated antigen 5\|Spag5** | 3.87246051 | 0.0435929 |
| **sulfatase 2\|Sulf2** | 3.87036674 | 0.00441282 |
| **histone deacetylase 4\|Hdac4** | 3.85937218 | 0.04062149 |
| **Sec61 beta subunit\|Sec61b** | 3.85499167 | 0.01652323 |
| **glucosamine-6-phosphate deaminase 1\|Gnpda1** | 3.85000202 | 0.00753885 |
| **coiled-coil domain containing 50\|Ccdc50** | 3.82823458 | 0.0079776 |
| **RNA binding motif protein 12 B2\|Rbm12b2** | 3.81735424 | 0.03665714 |
| **cell adhesion molecule 3\|Cadm3** | 3.80023202 | 0.04794045 |
| **filamin, alpha\|Flna** | 3.751354 | 0.01234048 |
| **vascular endothelial growth factor A\|Vegfa** | 3.73947505 | 0.01719525 |
| **family with sequence similarity 241, member A\|Fam241a** | 3.73026615 | 0.01020465 |
| **meningioma 1\|Mn1** | 3.7277615 | 0.01393496 |
| **tubulin, alpha 1A\|Tuba1a** | 3.70429969 | 0.0420074 |
| **RIKEN cDNA 2610028H24 gene\|2610028H24Rik** | 3.68899641 | 0.03994594 |
| **SMAD family member 7\|Smad7** | 3.66441526 | 0.04618995 |
| **cytoskeleton-associated protein 4\|Ckap4** | 3.63374454 | 0.00881584 |

| | | |
|---|---|---|
| **pleckstrin homology like domain, family A, member 1\|Phlda1** | 3.63367727 | 0.04467012 |
| **fibronectin type III domain containing 3B\|Fndc3b** | 3.61351086 | 0.01906245 |
| **frizzled class receptor 7\|Fzd7** | 3.59894267 | 0.02046178 |
| **procollagen-proline, 2-oxoglutarate 4-dioxygenase (proline 4-hydroxylase), alpha 1 polypeptide\|P4ha1** | 3.59691588 | 0.01435015 |
| **DENN/MADD domain containing 4A\|Dennd4a** | 3.58085196 | 0.01636425 |
| **cysteine-rich with EGF-like domains 2\|Creld2** | 3.58035091 | 0.00770529 |
| **WW domain binding protein 1 like\|Wbp1l** | 3.55513209 | 0.00533211 |
| **low density lipoprotein receptor\|Ldlr** | 3.54787694 | 0.04062149 |
| **xin actin-binding repeat containing 1\|Xirp1** | 3.5392545 | 0.01083837 |
| **myelin basic protein expression factor 2, repressor\|Myef2** | 3.53094085 | 0.03995983 |
| **folliculin\|Flcn** | 3.48247031 | 0.01210247 |
| **myocilin\|Myoc** | 3.45219983 | 0.01107458 |
| **DENN/MADD domain containing 5B\|Dennd5b** | 3.44874778 | 0.04029732 |
| **ubiquitination factor E4B\|Ube4b** | 3.4112675 | 0.01336038 |
| **ankyrin repeat domain 2 (stretch responsive muscle)\|Ankrd2** | 3.38367226 | 0.01312 |
| **PWWP domain containing 2B\|Pwwp2b** | 3.38161163 | 0.02232246 |
| **actin-binding LIM protein 1\|Ablim1** | 3.36105424 | 0.00524217 |
| **PCF11 cleavage and polyadenylation factor subunit\|Pcf11** | 3.35323871 | 0.02288639 |
| **growth arrest-specific 2 like 1\|Gas2l1** | 3.33348816 | 0.01393496 |
| **alanyl (membrane) aminopeptidase\|Anpep** | 3.32499391 | 0.03937018 |
| **ring finger protein 150\|Rnf150** | 3.29262801 | 0.02199621 |
| **paralemmin A kinase anchor protein\|Pakap** | 3.26504545 | 0.03982991 |
| **ATPase, class I, type 8B, member 2\|Atp8b2** | 3.26459174 | 0.01831478 |
| **Ras association (RalGDS/AF-6) and pleckstrin homology domains 1\|Raph1** | 3.26324435 | 0.01188289 |
| **sulfatase 1\|Sulf1** | 3.24460692 | 0.03908268 |
| **cystinosis, nephropathic\|Ctns** | 3.22788074 | 0.04104955 |
| **solute carrier family 22 (organic cation transporter), member 4\|Slc22a4** | 3.20973651 | 0.04144853 |
| **dishevelled segment polarity protein 2\|Dvl2** | 3.19815221 | 0.02476066 |
| **ER degradation enhancer, mannosidase alpha-like 1\|Edem1** | 3.1872383 | 0.04305617 |
| **cytokine receptor-like factor 1\|Crlf1** | 3.18292805 | 0.04259879 |
| **solute carrier family 22, member 23\|Slc22a23** | 3.06904838 | 0.02289676 |
| **tubulin, beta 5 class I\|Tubb5** | 3.05815577 | 0.03816213 |
| **HECT domain E3 ubiquitin protein ligase 2\|Hectd2** | 3.05098326 | 0.03809017 |
| **serine carboxypeptidase 1\|Scpep1** | 3.03478636 | 0.0420074 |
| **mesencephalic astrocyte-derived neurotrophic factor\|-** | 3.01255313 | 0.03858238 |
| **RUN and SH3 domain containing 2\|Rusc2** | 2.96292475 | 0.02123101 |
| **dual specificity phosphatase 1\|Dusp1** | 2.95055223 | 0.01411056 |
| **RIKEN cDNA 6430548M08 gene\|6430548M08Rik** | 2.94433773 | 0.03373278 |
| **mannosidase 2, alpha B2\|Man2b2** | 2.93409024 | 0.0484129 |
| **Expression decreased in β-DKO vs fl/fl control** | | |
| **UDP-Gal:betaGlcNAc beta 1,3-galactosyltransferase, polypeptide 2\|B3galt2** | 11.4338965 | 2.7188E-06 |
| **zinc finger protein 750\|Zfp750** | 9.13970537 | 0.00012572 |
| **aldehyde dehydrogenase 1 family, member L1\|Aldh1l1** | 8.41543806 | 0.00128174 |

| | | |
|---|---|---|
| **leucine rich repeat containing 38\|Lrrc38** | 8.17258487 | 1.8005E-05 |
| **MEF2 activating motif and SAP domain containing transcriptional regulator\|Mamstr** | 7.75273912 | 2.4859E-07 |
| **glycine N-methyltransferase\|Gnmt** | 7.51519006 | 0.00534687 |
| **glutathione S-transferase, theta 2\|Gstt2** | 7.25014622 | 0.03736185 |
| **perilipin 5\|Plin5** | 6.15304681 | 0.00274157 |
| **zinc finger protein 879\|Zfp879** | 6.13355465 | 0.00362567 |
| **predicted gene 4841\|Gm4841** | 6.01736804 | 0.01680409 |
| **polo like kinase 2\|Plk2** | 5.78643543 | 0.00657922 |
| **parathyroid hormone 1 receptor\|Pth1r** | 5.6312026 | 0.00506048 |
| **Cbp/p300-interacting transactivator, with Glu/Asp-rich carboxy-terminal domain, 4\|Cited4** | 5.59094959 | 0.01287261 |
| **ring finger protein 122\|Rnf122** | 5.32658923 | 0.01983834 |
| **CECR2, histone acetyl-lysine reader\|Cecr2** | 5.22087551 | 0.03469879 |
| **apolipoprotein L 10B\|Apol10b** | 5.03866065 | 0.01812929 |
| **G0/G1 switch gene 2\|G0s2** | 5.03002457 | 0.02808871 |
| **interferon-induced protein with tetratricopeptide repeats 3B\|Ifit3b** | 5.00253492 | 0.01469158 |
| **solute carrier family 40 (iron-regulated transporter), member 1\|Slc40a1** | 4.94374336 | 0.00614409 |
| **WSC domain containing 1\|Wscd1** | 4.92391602 | 0.00425523 |
| **RIKEN cDNA 6430571L13 gene\|6430571L13Rik** | 4.91955229 | 0.03038222 |
| **Rho GTPase activating protein 18\|Arhgap18** | 4.8287183 | 0.00131644 |
| **4-hydroxyphenylpyruvate dioxygenase-like\|Hpdl** | 4.82478487 | 0.00204405 |
| **synuclein, alpha interacting protein (synphilin)\|Sncaip** | 4.73964006 | 0.0006381 |
| **myosin regulatory light chain interacting protein\|Mylip** | 4.5218652 | 0.00066092 |
| **radical S-adenosyl methionine domain containing 2\|Rsad2** | 4.49226092 | 0.03042766 |
| **tubulin, gamma 2\|Tubg2** | 4.46478956 | 0.029444 |
| **HAUS augmin-like complex, subunit 4\|Haus4** | 4.43162011 | 0.01133451 |
| **ectonucleotide pyrophosphatase/phosphodiesterase 3\|Enpp3** | 4.4313483 | 0.00890655 |
| **NIM1 serine/threonine protein kinase\|Nim1k** | 4.39256762 | 0.04029732 |
| **acyl-CoA thioesterase 2\|Acot2** | 4.38582924 | 0.02864561 |
| **NAD(P)H dehydrogenase, quinone 1\|Nqo1** | 4.38159969 | 0.01192778 |
| **armadillo-like helical domain containing 4\|Armh4** | 4.3404354 | 0.00437052 |
| **DNA-damage-inducible transcript 4-like\|Ddit4l** | 4.31098581 | 0.03593761 |
| **dihydrodiol dehydrogenase (dimeric)\|Dhdh** | 4.28364601 | 0.00607362 |
| **zinc finger protein 52\|Zfp52** | 4.25352579 | 0.03756582 |
| **cortexin 3\|Ctxn3** | 4.21324324 | 0.01652323 |
| **indolethylamine N-methyltransferase\|Inmt** | 4.20640946 | 0.0072803 |
| **myogenic differentiation 1\|Myod1** | 4.20079844 | 0.01985801 |
| **Cbp/p300-interacting transactivator, with Glu/Asp-rich carboxy-terminal domain, 2\|Cited2** | 4.13755896 | 0.00232909 |
| **adhesion molecule with Ig like domain 3\|Amigo3** | 4.09394199 | 0.04104955 |
| **dual specificity phosphatase 10\|Dusp10** | 4.00580129 | 0.00066092 |
| **nuclear receptor subfamily 1, group D, member 1\|Nr1d1** | 3.95636641 | 0.02012522 |
| **thyroid hormone responsive\|Thrsp** | 3.84782958 | 0.01079525 |
| **carbonic anhydrase 14\|Car14** | 3.84019653 | 0.0044909 |
| **6-phosphofructo-2-kinase/fructose-2,6-biphosphatase 1\|Pfkfb1** | 3.76002189 | 0.0079776 |

| | | |
|---|---|---|
| **solute carrier family 26, member 6\|Slc26a6** | 3.75318929 | 0.00753885 |
| **kelch-like 33\|Klhl33** | 3.72979227 | 0.00216916 |
| **secretin receptor\|Sctr** | 3.72906345 | 0.02466707 |
| **MDS1 and EVI1 complex locus\|Mecom** | 3.64528803 | 0.03756582 |
| **carbonic anhydrase 4\|Car4** | 3.61749523 | 0.03181727 |
| **nuclear receptor subfamily 1, group H, member 3\|Nr1h3** | 3.58793363 | 0.01335667 |
| **microsomal glutathione S-transferase 1\|Mgst1** | 3.58431569 | 0.00200937 |
| **plexin domain containing 1\|Plxdc1** | 3.57848845 | 0.02704906 |
| **target of EGR1, member 1 (nuclear)\|Toe1** | 3.55096244 | 0.03038222 |
| **fibroblast growth factor receptor 4\|Fgfr4** | 3.38389371 | 0.02507614 |
| **septin 4\|Septin4** | 3.36587627 | 0.00998991 |
| **transmembrane and coiled-coil domains 4\|Tmco4** | 3.3215674 | 0.01204016 |
| **signal transducer and activator of transcription 5A\|Stat5a** | 3.28305043 | 0.02320622 |
| **pyruvate dehydrogenase phosphatase catalytic subunit 1\|Pdp1** | 3.21620341 | 0.01345282 |
| **phospholipase C, delta 4\|Plcd4** | 3.19970345 | 0.02442613 |
| **glutathione S-transferase, mu 2\|Gstm2** | 3.11890944 | 0.01723242 |
| **isocitrate dehydrogenase 1 (NADP+), soluble\|Idh1** | 3.10151072 | 0.02392418 |
| **polyamine oxidase (exo-N4-amino)\|Paox** | 3.04228349 | 0.04501694 |
| **argininosuccinate lyase\|Asl** | 3.01638018 | 0.04314435 |
| **pleckstrin homology domain containing, family H (with MyTH4 domain) member 3\|Plekhh3** | 2.99715921 | 0.01572652 |
| **sema domain, transmembrane domain (TM), and cytoplasmic domain, (semaphorin) 6C\|Sema6c** | 2.96476672 | 0.03276821 |
| **ankyrin repeat domain 9\|Ankrd9** | 2.89643711 | 0.03042766 |
| **potassium inwardly-rectifying channel, subfamily J, member 8\|Kcnj8** | 2.81555144 | 0.03756582 |
| **kyphoscoliosis peptidase\|Ky** | 2.78095136 | 0.04446621 |
| **ral guanine nucleotide dissociation stimulator\|Ralgds** | 2.73816817 | 0.04191859 |
| **CDC42 effector protein (Rho GTPase binding) 3\|Cdc42ep3** | 2.69824731 | 0.04218048 |
| **FAST kinase domains 1\|Fastkd1** | 2.65244329 | 0.04144358 |
| **caseinolytic mitochondrial matrix peptidase chaperone subunit\|Clpx** | 2.64991331 | 0.02898906 |

**Supplementary Table 3: Transcripts differentially expressed in EDL muscle between β-DKO mice treated with PBS and insulin.**

| | Fold change | False discovery rate |
|---|---|---|
| **Expression increased in β-DKO Ins vs β-DKO PBS** | | |
| **lactate dehydrogenase B\|Ldhb** | 4.353275 | 0.080173 |
| **microtubule-associated protein 1 A\|Map1a** | 4.070883 | 0.055235 |
| **EPS8-like 2\|Eps8l2** | 3.439704 | 0.003969 |
| **myoglobin\|Mb** | 2.250541 | 0.000735 |
| **creatine kinase, mitochondrial 2\|Ckmt2** | 1.961628 | 0.011665 |
| **myosin binding protein C, slow-type\|Mybpc1** | 1.599328 | 0.084026 |
| **myosin, heavy polypeptide 1, skeletal muscle, adult\|Myh1** | 1.594162 | 0.009398 |
| **mitochondrially encoded NADH dehydrogenase 2\|ND2** | 1.396794 | 0.017172 |
| **mitochondrially encoded cytochrome c oxidase III\|COX3** | 1.197349 | 0.09742 |
| **Expression decreased in β-DKO Ins vs β-DKO PBS** | | |
| **aldolase A, fructose-bisphosphate\|Aldoa** | 1.209698 | 0.09742 |
| **ryanodine receptor 1, skeletal muscle\|Ryr1** | 1.287901 | 0.055235 |
| **myosin binding protein C, fast-type\|Mybpc2** | 1.417734 | 0.008806 |
| **dual specificity phosphatase 1\|Dusp1** | 1.439174 | 0.080173 |
| **tropomyosin 1, alpha\|Tpm1** | 1.511536 | 9.97E-07 |
| **myosin, heavy polypeptide 4, skeletal muscle\|Myh4** | 1.697096 | 4.00E-21 |
| **frizzled class receptor 7\|Fzd7** | 1.983316 | 0.000208 |
| **tectonin beta-propeller repeat containing 2\|Tecpr2** | 2.165953 | 0.007169 |
| **potassium voltage-gated channel, subfamily G, member 4\|Kcng4** | 2.625393 | 0.010772 |
| **potassium voltage-gated channel, subfamily F, member 1\|Kcnf1** | 2.877588 | 0.015407 |
| **upstream transcription factor family member 3\|Usf3** | 2.961892 | 5.16E-09 |
| **RPTOR independent companion of MTOR, complex 2\|Rictor** | 3.522655 | 1.16E-05 |
| **cadherin 4\|Cdh4** | 4.737905 | 3.12E-05 |
| **Myosin regulatory light chain 12B\|-** | 13.11259 | 0.056071 |
| **aldolase A, fructose-bisphosphate\|Aldoa** | 1.209698 | 0.09742 |
| **ryanodine receptor 1, skeletal muscle\|Ryr1** | 1.287901 | 0.055235 |
| **myosin binding protein C, fast-type\|Mybpc2** | 1.417734 | 0.008806 |
| **dual specificity phosphatase 1\|Dusp1** | 1.439174 | 0.080173 |
| **tropomyosin 1, alpha\|Tpm1** | 1.511536 | 9.97E-07 |
| **myosin, heavy polypeptide 4, skeletal muscle\|Myh4** | 1.697096 | 4.00E-21 |
| **frizzled class receptor 7\|Fzd7** | 1.983316 | 0.000208 |
| **tectonin beta-propeller repeat containing 2\|Tecpr2** | 2.165953 | 0.007169 |
| **potassium voltage-gated channel, subfamily G, member 4\|Kcng4** | 2.625393 | 0.010772 |
| **potassium voltage-gated channel, subfamily F, member 1\|Kcnf1** | 2.877588 | 0.015407 |
| **upstream transcription factor family member 3\|Usf3** | 2.961892 | 5.16E-09 |
| **RPTOR independent companion of MTOR, complex 2\|Rictor** | 3.522655 | 1.16E-05 |
| **cadherin 4\|Cdh4** | 4.737905 | 3.12E-05 |

**Supplementary Table 4: Transcripts differentially expressed in soleus muscle between β-DKO mice treated with PBS and insulin.**

| | Fold change | False discovery rate |
|---|---|---|
| **Expression increased in β-DKO Ins vs β-DKO PBS** | | |
| **B cell leukemia/lymphoma 3\|Bcl3** | 0.08841787 | 0.02524168 |
| **ankyrin repeat domain 1 (cardiac muscle)\|-** | 0.09930934 | 0.00164911 |
| **predicted gene 13889\|Gm13889** | 0.11185993 | 2.6666E-10 |
| **hemoglobin alpha, adult chain 1\|Hba-a1** | 0.12352729 | 0.02045674 |
| **hemoglobin, beta adult t chain\|Hbb-bt** | 0.17792673 | 0.03952168 |
| **ankyrin repeat domain 35\|Ankrd35** | 0.17805746 | 0.00783436 |
| **leucine rich repeat containing 38\|Lrrc38** | 0.18410922 | 0.00013545 |
| **CD68 antigen\|Cd68** | 0.18857368 | 0.03952168 |
| **heat shock protein 1B\|Hspa1b** | 0.18861857 | 0.00036588 |
| **potassium voltage-gated channel, subfamily G, member 2\|Kcng2** | 0.23013122 | 0.04990613 |
| **cytochrome P450, family 39, subfamily a, polypeptide 1\|Cyp39a1** | 0.24787265 | 0.01905892 |
| **phosphodiesterase 4B, cAMP specific\|Pde4b** | 0.26250656 | 0.01421318 |
| **heat shock protein 1\|Hspb1** | 0.26311706 | 0.00732982 |
| **H4 clustered histone 9\|H4c9** | 0.2717768 | 0.01890276 |
| **dual specificity phosphatase 10\|Dusp10** | 0.27977918 | 0.01619584 |
| **predicted gene 3893\|-** | 0.28164255 | 0.03952168 |
| **heat shock protein 1A\|Hspa1a** | 0.31126719 | 0.00192608 |
| **Expression increased in β-DKO Ins vs β-DKO PBS** | | |
| **phosphoenolpyruvate carboxykinase 1, cytosolic\|Pck1** | 27.9690394 | 0.03538523 |
| **dachshund family transcription factor 1\|Dach1** | 14.9714951 | 1.7965E-11 |
| **predicted gene 20547\|-** | 8.9544479 | 0.00141646 |
| **pentraxin 4\|Ptx4** | 8.48018996 | 2.8531E-06 |
| **adenomatosis polyposis coli down-regulated 1\|-** | 8.32195377 | 0.00471186 |
| **phosphodiesterase 3B, cGMP-inhibited\|Pde3b** | 8.107231 | 0.00642456 |
| **solute carrier family 4 (anion exchanger), member 4\|Slc4a4** | 8.09514442 | 0.01391817 |
| **NCK-associated protein 5\|Nckap5** | 7.91214405 | 0.01378667 |
| **ADAMTS-like 1\|Adamtsl1** | 7.88757225 | 0.00642456 |
| **HECT, C2 and WW domain containing E3 ubiquitin protein ligase 2\|Hecw2** | 6.66513645 | 0.00471186 |
| **phospholipase C, beta 1\|Plcb1** | 6.40426037 | 0.000363 |
| **acyl-CoA thioesterase 1\|Acot1** | 6.23687792 | 0.01064114 |
| **family with sequence similarity 117, member B\|Fam117b** | 5.43000925 | 0.01504768 |
| **flavin containing monooxygenase 2\|Fmo2** | 5.24272618 | 0.000363 |
| **RIKEN cDNA 9930111J21 gene 1\|9930111J21Rik1** | 5.12557903 | 0.01382766 |
| **fibronectin 1\|Fn1** | 4.99108511 | 0.00037872 |
| **butyrophilin-like 9\|Btnl9** | 4.83043807 | 0.00642456 |
| **C1q and tumor necrosis factor related protein 1\|C1qtnf1** | 4.77202699 | 0.04353263 |
| **reelin\|Reln** | 4.74882176 | 0.00192608 |
| **ajuba LIM protein\|Ajuba** | 4.46017912 | 0.00960412 |

| | | |
|---|---|---|
| **predicted gene, 26566\|-** | 4.424546 | 0.01582583 |
| **myelin basic protein expression factor 2, repressor\|Myef2** | 4.29026669 | 0.00642456 |
| **DEPP1 autophagy regulator\|Depp1** | 4.18411514 | 0.00192608 |
| **Kruppel-like factor 13\|Klf13** | 4.18039964 | 0.01150915 |
| **guanine deaminase\|Gda** | 4.14941667 | 0.00564087 |
| **Rap guanine nucleotide exchange factor (GEF) 5\|Rapgef5** | 3.99789603 | 0.03043476 |
| **ATPase, Ca++ transporting, plasma membrane 4\|Atp2b4** | 3.89576952 | 0.00471186 |
| **guanylate binding protein 7\|Gbp7** | 3.81208348 | 0.02371283 |
| **angiomotin-like 2\|Amotl2** | 3.72335557 | 0.00783436 |
| **papilin, proteoglycan-like sulfated glycoprotein\|Papln** | 3.62550159 | 0.00643268 |
| **protein tyrosine phosphatase, receptor type, B\|Ptprb** | 3.61551009 | 0.00036588 |
| **MAM domain containing 2\|Mamdc2** | 3.61063068 | 0.01796555 |
| **cellular communication network factor 3\|Ccn3** | 3.60055621 | 0.04016193 |
| **ATP-binding cassette, sub-family B (MDR/TAP), member 1A\|Abcb1a** | 3.593384 | 0.00164911 |
| **bone morphogenetic protein 6\|-** | 3.55786969 | 0.017957 |
| **junctional cadherin 5 associated\|Jcad** | 3.55134259 | 0.00037872 |
| **slit guidance ligand 3\|Slit3** | 3.46589552 | 0.00643268 |
| **tissue inhibitor of metalloproteinase 3\|Timp3** | 3.44971659 | 0.01382766 |
| **golgi associated kinase 1B\|Gask1b** | 3.44087829 | 0.00564087 |
| **phosphatidylinositol-3,4,5-trisphosphate-dependent Rac exchange factor 2\|Prex2** | 3.43321303 | 0.01796555 |
| **LIF receptor alpha\|Lifr** | 3.42774368 | 0.04353263 |
| **forkhead box K1\|Foxk1** | 3.4257517 | 0.03952168 |
| **shroom family member 4\|Shroom4** | 3.27099122 | 0.02606346 |
| **ATP-binding cassette, sub-family A (ABC1), member 5\|Abca5** | 3.24054358 | 0.00564087 |
| **mcf.2 transforming sequence-like\|Mcf2l** | 3.17635601 | 0.00642456 |
| **folliculin interacting protein 2\|Fnip2** | 3.17305993 | 0.03043476 |
| **zinc finger and BTB domain containing 10\|Zbtb10** | 3.08119871 | 0.03952168 |
| **adhesion G protein-coupled receptor F5\|Adgrf5** | 3.05350087 | 0.03538523 |
| **jagged 1\|Jag1** | 3.02933175 | 0.03952168 |
| **SMAD family member 7\|Smad7** | 3.00268082 | 0.03952168 |
| **early B cell factor 1\|Ebf1** | 2.99380555 | 0.03952168 |
| **integrin alpha 6\|Itga6** | 2.94607614 | 0.017957 |
| **plexin A2\|Plxna2** | 2.93543586 | 0.0437692 |
| **caveolae associated 2\|Cavin2** | 2.85356232 | 0.00732982 |
| **Von Willebrand factor\|Vwf** | 2.58427128 | 0.04105607 |
| **proline rich 13\|Prr13** | 0.36175919 | 0.04505914 |
| **rhomboid like 1\|Rhbdl1** | 0.32862046 | 0.00642456 |
| **tenascin C\|Tnc** | 0.32746827 | 0.02732075 |

**Supplementary Table 5: Gene Ontology biological process results based on transcripts differentially expressed in EDL muscle between control fl/fl mice and β-DKO mice obtained from the Mouse Portal of the Rate Genome Database.**

| biological process | p value |
|---|---|
| muscle contraction | 7.31E-12 |
| muscle system process | 3.20E-11 |
| voluntary skeletal muscle contraction | 4.53E-09 |
| twitch skeletal muscle contraction | 4.53E-09 |
| striated muscle contraction | 5.01E-08 |
| slow-twitch skeletal muscle fiber contraction | 9.39E-08 |
| skeletal muscle contraction | 3.36E-07 |
| multicellular organismal movement | 1.46E-06 |
| musculoskeletal movement | 1.46E-06 |
| transition between fast and slow fiber | 2.77E-06 |
| regulation of the force of skeletal muscle contraction | 8.34E-06 |
| regulation of skeletal muscle contraction by chemo-mechanical energy conversion | 8.34E-06 |
| regulation of skeletal muscle adaptation | 1.27E-05 |

**Supplementary Table 6: Gene Ontology biological process results based on transcripts differentially expressed in soleus muscle between control fl/fl mice and β-DKO mice obtained from the Mouse Portal of the Rate Genome Database.**

| biological process | p value |
|---|---|
| tissue development | 4.21E-23 |
| anatomical structure development | 1.39E-21 |
| multicellular organism development | 1.14E-20 |
| extracellular matrix organization | 1.82E-20 |
| developmental process | 1.85E-20 |
| external encapsulating structure organization | 2.26E-20 |
| extracellular structure organization | 2.26E-20 |
| system development | 2.31E-20 |
| animal organ development | 4.92E-19 |
| collagen fibril organization | 5.35E-18 |
| positive regulation of cellular process | 5.00E-16 |
| positive regulation of biological process | 5.26E-16 |
| anatomical structure morphogenesis | 3.56E-15 |
| circulatory system development | 4.78E-15 |
| regulation of locomotion | 5.21E-15 |
| regulation of cell migration | 5.24E-15 |
| regulation of cell motility | 1.32E-14 |
| cell differentiation | 2.06E-14 |
| cellular developmental process | 2.82E-14 |
| regulation of multicellular organismal process | 3.40E-14 |
| regulation of programmed cell death | 1.18E-13 |
| negative regulation of cellular process | 1.56E-13 |
| regulation of developmental process | 2.43E-13 |
| cell adhesion | 5.88E-13 |
| regulation of apoptotic process | 7.30E-13 |
| negative regulation of biological process | 8.39E-13 |
| cellular response to chemical stimulus | 8.51E-13 |
| regulation of cell differentiation | 1.35E-12 |
| skeletal system development | 3.71E-12 |
| response to organic substance | 4.60E-12 |
| regulation of cell population proliferation | 6.69E-12 |
| vasculature development | 2.03E-11 |
| tube development | 5.58E-11 |
| response to external stimulus | 8.56E-11 |
| blood vessel development | 8.59E-11 |
| supramolecular fiber organization | 1.03E-10 |
| regulation of response to stimulus | 1.29E-10 |
| regulation of cell adhesion | 1.40E-10 |
| cellular response to oxygen-containing compound | 2.33E-10 |
| cell development | 2.85E-10 |

| | |
|---|---|
| biological regulation | 3.26E-10 |
| regulation of cellular process | 7.32E-10 |
| response to chemical | 7.38E-10 |
| tube morphogenesis | 7.72E-10 |
| homeostatic process | 7.76E-10 |
| positive regulation of cell migration | 8.40E-10 |
| cellular response to organic substance | 1.02E-09 |
| regulation of biological process | 1.18E-09 |
| response to acid chemical | 1.28E-09 |
| negative regulation of programmed cell death | 1.98E-09 |
| positive regulation of cell motility | 2.46E-09 |
| multicellular organismal-level homeostasis | 2.67E-09 |
| multicellular organismal process | 2.78E-09 |
| negative regulation of developmental process | 2.82E-09 |
| response to cytokine | 2.96E-09 |
| connective tissue development | 3.00E-09 |
| positive regulation of locomotion | 4.52E-09 |
| regulation of metabolic process | 5.94E-09 |
| ossification | 6.55E-09 |
| negative regulation of apoptotic process | 7.14E-09 |
| cell population proliferation | 9.51E-09 |
| cell surface receptor signaling pathway | 1.11E-08 |
| heart development | 1.20E-08 |
| positive regulation of metabolic process | 1.26E-08 |
| response to oxygen-containing compound | 1.43E-08 |
| anatomical structure formation involved in morphogenesis | 1.77E-08 |
| epithelium development | 1.98E-08 |
| response to endogenous stimulus | 2.34E-08 |
| cell migration | 2.60E-08 |
| sensory perception of chemical stimulus | 2.63E-08 |
| positive regulation of programmed cell death | 2.99E-08 |
| regulation of multicellular organismal development | 3.38E-08 |
| response to organonitrogen compound | 3.68E-08 |
| cartilage development | 3.70E-08 |
| cellular response to acid chemical | 4.84E-08 |
| nervous system development | 5.80E-08 |
| response to lipid | 5.87E-08 |
| regulation of response to external stimulus | 6.24E-08 |
| regulation of signaling | 6.40E-08 |
| negative regulation of multicellular organismal process | 6.99E-08 |
| enzyme-linked receptor protein signaling pathway | 7.67E-08 |
| regulation of signal transduction | 8.23E-08 |
| cellular response to cytokine stimulus | 8.85E-08 |
| blood vessel morphogenesis | 1.03E-07 |
| animal organ morphogenesis | 1.22E-07 |
| regulation of primary metabolic process | 1.24E-07 |

| | |
|---|---|
| sensory perception of smell | 1.29E-07 |
| positive regulation of apoptotic process | 1.36E-07 |
| positive regulation of nitrogen compound metabolic process | 1.54E-07 |
| muscle tissue development | 1.58E-07 |
| cellular process | 1.76E-07 |
| regulation of cellular component organization | 1.82E-07 |
| positive regulation of cell adhesion | 2.16E-07 |
| regulation of cellular metabolic process | 2.27E-07 |
| negative regulation of cell differentiation | 2.35E-07 |
| positive regulation of cellular metabolic process | 2.53E-07 |
| bone development | 2.72E-07 |
| response to inorganic substance | 2.73E-07 |
| positive regulation of cell differentiation | 2.74E-07 |
| regulation of cell communication | 3.16E-07 |
| regulation of epithelial cell proliferation | 3.25E-07 |
| cellular component organization | 3.66E-07 |
| response to nitrogen compound | 4.08E-07 |
| cell motility | 4.11E-07 |
| response to amino acid | 4.81E-07 |
| cellular response to stimulus | 5.37E-07 |
| skeletal system morphogenesis | 5.73E-07 |
| regulation of cell-substrate adhesion | 5.86E-07 |
| negative regulation of immune system process | 5.93E-07 |
| positive regulation of macromolecule metabolic process | 7.22E-07 |
| positive regulation of cell population proliferation | 8.29E-07 |
| positive regulation of cellular component organization | 8.35E-07 |
| chondrocyte differentiation | 8.85E-07 |
| embryo development | 8.91E-07 |
| regulation of cell development | 9.34E-07 |
| positive regulation of multicellular organismal process | 1.00E-06 |
| cell-substrate adhesion | 1.04E-06 |
| renal system development | 1.22E-06 |
| positive regulation of biosynthetic process | 1.35E-06 |
| positive regulation of cell-substrate adhesion | 1.38E-06 |
| regulation of macromolecule metabolic process | 1.48E-06 |
| apoptotic process | 1.51E-06 |
| regulation of substrate adhesion-dependent cell spreading | 1.63E-06 |
| positive regulation of developmental process | 1.63E-06 |
| regulation of cell activation | 1.74E-06 |
| regulation of immune system process | 2.05E-06 |
| regulation of fat cell differentiation | 2.12E-06 |
| positive regulation of cellular biosynthetic process | 2.14E-06 |
| regulation of phosphate metabolic process | 2.18E-06 |
| regulation of phosphorus metabolic process | 2.20E-06 |
| regulation of nitrogen compound metabolic process | 2.25E-06 |
| negative regulation of cell adhesion | 2.31E-06 |

| | |
|---|---|
| kidney development | 2.31E-06 |
| regulation of myotube differentiation | 2.33E-06 |
| regulation of biosynthetic process | 2.56E-06 |
| positive regulation of signal transduction | 2.61E-06 |
| regulation of smooth muscle cell migration | 2.66E-06 |
| negative regulation of cell population proliferation | 2.85E-06 |
| cellular component organization or biogenesis | 3.34E-06 |
| biomineral tissue development | 3.42E-06 |
| skeletal muscle cell differentiation | 3.71E-06 |
| positive regulation of smooth muscle cell migration | 3.71E-06 |
| positive regulation of signaling | 3.82E-06 |
| response to stress | 3.85E-06 |
| response to stimulus | 3.96E-06 |
| regulation of gene expression | 4.34E-06 |
| epithelial cell proliferation | 4.94E-06 |
| response to hormone | 5.07E-06 |
| cellular response to amino acid stimulus | 5.14E-06 |
| regulation of cellular biosynthetic process | 5.28E-06 |
| transmembrane receptor protein tyrosine kinase signaling pathway | 5.65E-06 |
| programmed cell death | 5.77E-06 |
| cell death | 5.91E-06 |
| sensory perception | 6.22E-06 |
| positive regulation of cell communication | 6.23E-06 |
| regulation of phosphorylation | 6.56E-06 |
| positive regulation of response to stimulus | 6.59E-06 |
| negative regulation of response to stimulus | 8.72E-06 |
| regulation of small molecule metabolic process | 8.79E-06 |
| positive regulation of DNA-templated transcription | 9.51E-06 |
| regulation of intracellular signal transduction | 9.84E-06 |
| positive regulation of RNA biosynthetic process | 1.00E-05 |
| endothelium development | 1.01E-05 |
| bone trabecula morphogenesis | 1.03E-05 |
| positive regulation of transcription by RNA polymerase II | 1.16E-05 |
| collagen metabolic process | 1.22E-05 |
| negative regulation of locomotion | 1.27E-05 |
| regulation of catalytic activity | 1.27E-05 |
| tendon development | 1.30E-05 |
| regulation of cold-induced thermogenesis | 1.34E-05 |
| tissue morphogenesis | 1.36E-05 |
| muscle structure development | 1.40E-05 |
| positive regulation of epithelial cell proliferation | 1.49E-05 |
| response to molecule of bacterial origin | 1.51E-05 |
| regulation of extracellular matrix organization | 1.69E-05 |
| regulation of molecular function | 1.76E-05 |
| bone mineralization | 1.88E-05 |
| endothelial cell differentiation | 1.97E-05 |

| | |
|---|---|
| chondrocyte development | 2.20E-05 |
| cellular response to lipid | 2.26E-05 |
| homeostasis of number of cells | 2.37E-05 |
| response to lipopolysaccharide | 2.42E-05 |
| intracellular signal transduction | 2.42E-05 |
| angiogenesis | 2.55E-05 |
| positive regulation of protein metabolic process | 2.75E-05 |
| endochondral bone morphogenesis | 3.10E-05 |
| regeneration | 3.17E-05 |
| response to abiotic stimulus | 3.28E-05 |
| response to wounding | 3.41E-05 |
| positive regulation of cell projection organization | 3.42E-05 |
| regulation of biological quality | 3.56E-05 |
| positive regulation of smooth muscle cell proliferation | 3.64E-05 |
| locomotion | 3.77E-05 |
| regulation of neuron projection development | 4.06E-05 |
| wound healing | 4.15E-05 |
| regulation of body fluid levels | 4.27E-05 |
| negative regulation of cell motility | 4.43E-05 |
| cellular response to endogenous stimulus | 4.44E-05 |
| endothelial cell development | 4.48E-05 |
| regulation of anatomical structure size | 4.88E-05 |
| trabecula morphogenesis | 5.00E-05 |
| chemotaxis | 5.12E-05 |
| positive regulation of RNA metabolic process | 5.21E-05 |
| taxis | 5.41E-05 |
| response to extracellular stimulus | 5.69E-05 |
| negative regulation of leukocyte activation | 5.86E-05 |
| artery morphogenesis | 5.88E-05 |
| bone trabecula formation | 5.89E-05 |
| positive regulation of cell development | 5.98E-05 |
| negative regulation of metabolic process | 6.02E-05 |
| cellular response to oxidative stress | 6.43E-05 |
| regulation of inflammatory response | 6.75E-05 |
| positive regulation of macromolecule biosynthetic process | 7.09E-05 |
| regulation of anatomical structure morphogenesis | 7.17E-05 |
| muscle organ development | 7.41E-05 |
| regulation of leukocyte activation | 7.46E-05 |
| tissue remodeling | 7.47E-05 |
| positive regulation of catalytic activity | 8.24E-05 |
| regulation of DNA-templated transcription | 9.03E-05 |
| negative regulation of cell migration | 9.20E-05 |
| neutrophil homeostasis | 9.21E-05 |
| response to oxidative stress | 9.23E-05 |
| neurogenesis | 9.34E-05 |
| regulation of RNA metabolic process | 9.42E-05 |

| | |
|---|---|
| regulation of RNA biosynthetic process | 9.54E-05 |
